# Body-plan reorganization in a sponge correlates with microbiome change

**DOI:** 10.1101/2022.07.22.501172

**Authors:** Sergio Vargas, Laura Leiva, Michael Eitel, Franziska Curdt, Sven Rohde, Christopher Arnold, Michael Nickel, Peter Schupp, William D. Orsi, Maja Adamska, Gert Wörheide

## Abstract

Mounting evidence suggests that animals and their associated bacteria interact via intricate molecular mechanisms, and it is hypothesized that disturbances to the microbiome can influence animal development. Sponges diverged from other animals more than 750 MYA and represent one of the earliest branching animal phyla that exhibit symbiotic relationships with diverse bacteria. Over 41 microbial phyla have been found in association with sponges, forming a holobiont that is integral to aquatic ecosystems worldwide. Sponge-associated microbes contain an enriched set of proteins bearing eukaryotic-like domains, and their metabolism supports the host with nutrients. This indicates strong physiological interconnections in the holobiont, which are thought to be modulated by sponge immunity and pattern-recognition proteins. Despite the hypothesized tight physiological integration and ancient origin of the sponge holobiont, the effect of changes in the symbiotic community on the sponge metabolism and morphogenesis remains poorly understood. Here, we show that the loss of a key microbial sponge symbiont correlates with a stark body plan reorganization of the sponge host. This reorganization is coupled with broad transcriptomic changes and includes the modulation of signaling pathways known to be involved in morphogenesis and innate immune response in sponges and other animals. This study provides a combined genetic, physiological, and morphological assessment of the effect of changes in the microbiome on sponge post-embryonic development and homeostasis. The drastic microbiome reorganization and the correlated response observed in the sponge host provide evidence for a coupling between sponge transcriptomic state and the state of its microbiome. Our results suggest that sponges use molecular mechanisms to respond to changes in their microbiome and that the ability to sense and respond to microbiome perturbations has deep evolutionary origins among animals.

## Introduction

In different bilaterian animal model systems, the microbiome can alter numerous host physiological and developmental processes. The renewal of the intestinal epithelium in zebrafish, for instance, is co-regulated by signals from its microbiome (Cheesman, Neal, Mittge, Seredick, & Guillemin, 2011). Brain development in mice also appears to be modulated by the gut microbiota, and evidence exists for a role of the microbiome on mouse behavior (Heijtz et al., 2011; Sampson & Mazmanian, 2015; Sharon, Sampson, Geschwind, & Mazmanian, 2016). Non-bilaterian animals also appear to interact in complex ways with their microbiomes. In Cnidaria, likely the sister group to the Bilateria (Simion et al., 2017), associated bacteria modulate the contraction behavior of the polyps of *Hydra vulgaris* (Klimovich et al., 2017). Thus, host-microbiome interactions appear to have deep evolutionary roots in the animal lineage.

Sponges, one of the two Phyla most likely representing the sister group to all remaining animal lineages (King & Rokas, 2017), harbor bacterial communities of varying complexity and richness (Thomas et al., 2016) and form holobionts with different degrees of physiological integration (Webster & Thomas, 2016). For instance, sponges that possess cyanobacteria-dominated microbiomes (*i.e*., cyanosponges) can be mixotrophs combining filter-feeding with symbiont-derived nutrition (Hudspith et al., 2022) or autotrophs, in which case their photosynthetic symbionts can provide up to 80% of their carbon requirements (Usher, 2008). Field experiments have shown that in some cyanosponges, photosymbiont loss can lead to a 42% reduction in biomass after a two-week exposure to shading (Thacker, 2005) and that the sponge hosts do not compensate for the reduction in symbiont-derived carbon resulting from shading through increased filter-feeding (Freeman & Thacker, 2011). These observations indicate a high degree of physiological integration between cyanosponges and their microbiomes.

Also, field experiments on cyanobacteria-bearing keratose sponges have reported that these sponges change their morphology, developing oscular lobes or chimneys, only after losing their cyanobacterial symbionts upon transplantation to deeper (*e.g*., 200m) waters (Maldonado & Young, 1998). These changes have been interpreted as an adjustment of the sponges to increase their filtering capacity in response to a reduced symbiont-derived carbon input (Maldonado & Young, 1998). They also suggest that a compositional reorganization of the sponge microbiome in response to environmental changes can result in a post-embryonic developmental reorganization of the sponge body plan. To study this, we use the autotrophic cyanosponge *Lendenfeldia chondrodes*, a common salt-water aquarium sponge (Galitz et al., 2018). Under normal culture conditions, *L. chondrodres* displays two different colorations and growth forms depending on the exposure of the sponge tissues to light (Fig. 1). Sections of the sponge exposed to light are purple to dark blue and display a foliose growth form, while shaded parts of the sponge are white and grow to form thread-like projections (Fig. 1). Here, we investigate the effect of shading on *L. chondrodes*’ microbiome, host morphology and host transcriptional state. Shading *L. chondrodes* causes a compositional accommodation of its microbiome upon loss of its cyanobacterial photosymbionts and results in a marked host response. This response is characterized by a reorganization of the sponge body-plan and growth form, and it is accompanied by a broad transcriptional response involving immune and developmental coexpression modules. In conjunction, these results offer support for a mechanistic coupling between the sponge physiological and transcriptional state and its microbiome, allowing the host to respond to changes in their symbiotic communities triggered by the environment.

**Figure 1.**
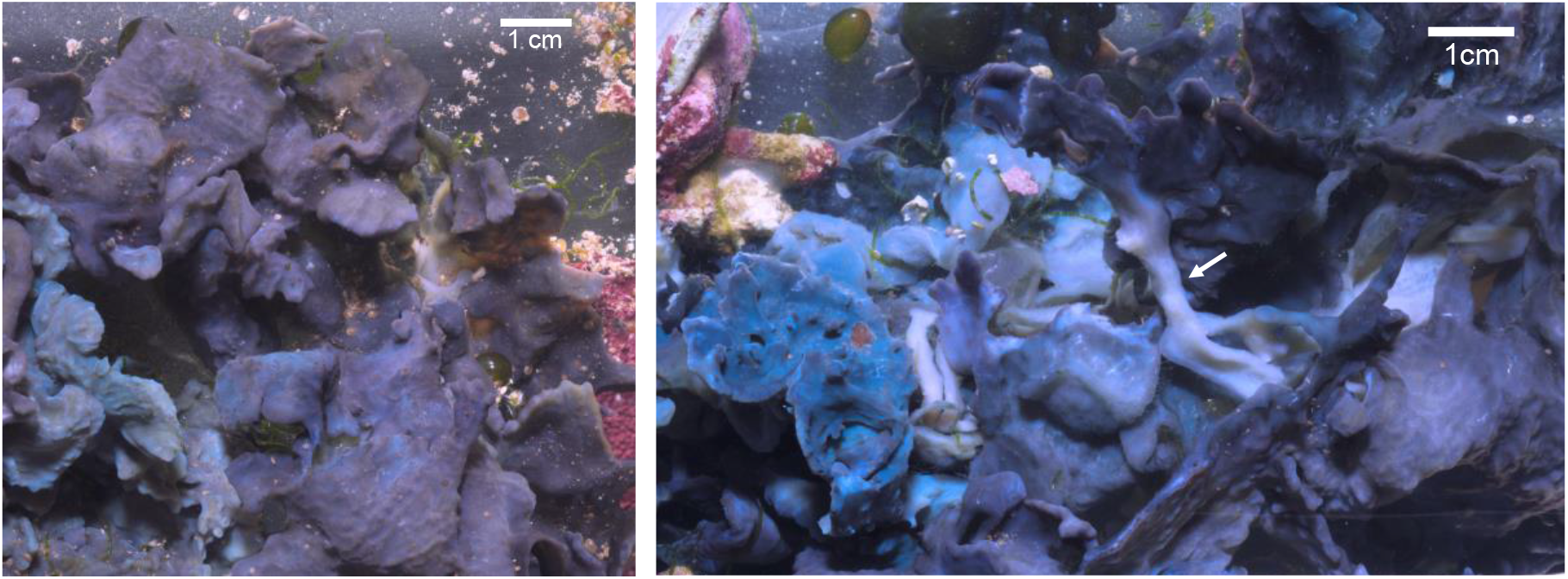
*Lendenfeldia chondrodes* growing under control conditions. Left: sponge viewed from the top, note the flattened, foliose morphology. Right: Bottom of the sponge shown in the left panel. White elongated sections can be seen (white left arrow) connecting foliose sections of the sponge.

## Methods and Methods

### Description of the aquarium system

Animals are kept in a 642 L marine aquarium under a 12h day, 12h night cycle and controlled temperature and pH. Based on hourly measurements over one year (2017), average water temperature and pH are 24.92 ± 0.24 °C and 8.30 ± 0.14. Based on weakly measurements over one year (2017), the average PO_4_^-3^, NO_2_^-^ and NO_3_^-^ concentrations are 0.092 ± 0.071 mg/L, 0.014 ± 0.072 mg/L, 2.681 ± 3.882 mg/L; the concentration of NH_3_/NH_4_^+^ was consistently below detection (*i.e*., < 0.05 mg/L). These values are monitored hourly and weekly and are stable. Thus, they represent baseline values for the system. All experiments were carried out in a tank connected to the main 642L aquarium tank through two water inlets and one central outlet that leads the water back to the system water reservoir from which is pumped back to the aquarium’s main and experimental tanks. Water flows constantly between the two systems so that sponge explants are exposed to the same temperature, pH, and nutrients measures in the main water tank. To avoid sediment accumulation on the experimental system, each inlet is connected to a sediment trap through which the water flows before entering the experimental area of the tank. The water in the experimental tank then flows to the main central water outlet. The experimental system was arranged in a symmetrical manner to avoid confounding factors like water flow. A graphical description of the aquarium and experimental system can be found in Suppl. Figure 1.

### Shading experiments

Explants (1cm in diameter) were exposed to shading in the same aquarium system where the parental sponges are kept. Shading was achieved using a black, plastic foil mounted on top of plexiglas panels covering the aquarium space where the explants were placed. Neither the plastic foil nor the plexiglass panels were in contact with the water or blocked the water movement, *i.e*., they were placed on top of the aquarium, allowing water to move in the system from the water inlets to the central outlet and only blocking the light.

Control and treatment explants were placed on (open) petri dishes, separated to avoid contact (and fusion) and were visually inspected every week. The petri dishes were manually cleaned using a Pasteur pipette if sand or fine sediment accumulated. This process was done using light and generally took less than one hour. The sponges were covered (if necessary) again after each check/cleaning. Due to the symmetrical arrangement of the water inflow in the experimental tank and the presence of a single, central outflow, both control and treatment sponges were exposed to similar conditions in terms of temperature, nutrients, water pH, dissolved oxygen, and exposure to water currents thus allowing us to eliminate potential confounding effects by these variables from the experiment. Treatment sponge explants were completely shaded by the black plastic foil or were allowed to receive only scattered light (*i.e*., Photosynthetic Active Radiation (PAR) levels below detection). This setup allowed us to constrain the possible confounding effect light deprivation may have on the sponges.

Sampling was done after at least 12 weeks (and no more than 15 weeks) of exposure to darkness using liquid nitrogen to flash freeze the sponges, which were rapidly transferred from the aquarium to 8mL vials filled with liquid nitrogen. To avoid transferring other organisms commonly found in the aquarium, like small amphipods or polychaetes, the sponge explants were visually inspected, carefully cleaned with a Pasteur pipette and transferred to the liquid nitrogen containing vial. Tissue samples for RNA extraction were kept at −80 °C. Other sponge samples were fixed in 96% ethanol and kept at −20 °C until further processing.

### 16S V4 amplicon sequencing and evaluation of community composition changes in response to bleaching

To characterize changes in the composition of *L. chondrodes*’ microbiome upon exposure to shading and loss of cyanobacterial symbionts, control and sponge explants exposed to shading (*i.e*., direct light was completely blocked or only scattered light was allowed to reach explants) for three, six, nine and twelve weeks were sampled, fixed in ethanol 96%, extracted using the Macherey-Nagel Nucleospin Tissue DNA extraction kit and used to amplify the V4 region of the 16S rRNA gene using indexed primers 515F and 806R (see Pichler et al., 2018). Amplicons were 150bp pair-end sequenced on an Illumina MiniSeq and the resulting reads were processed using *vsearch* (Rognes, Flouri, Nichols, Quince, & Mahé, 2016) to define Operational Taxonomic Units (OTU) using a cutoff distance of 97% similarity and to obtain a Sample by OTU abundance matrix. Community composition was analyzed in R (R Development Core Team, 2008) using the package vegan (Oksanen et al., 2017). We used an NMDS with the Bray-Curtis distance to visualize the similarity of the samples’ bacterial communities and an ANOSIM to test the hypothesis of no effect of the experimental treatments (*i.e*., direct shading *vs*. scattered light, and shading + scattered light *vs*. control condition) on the composition of the sponge microbiome. In addition, for the nine most abundant core OTUs (Vargas, Leiva, & Wörheide, 2021) we tested for significant changes in OUT frequency in control *vs*. shaded sponges using Hotelling’s T-Square test. This test was followed by individual, Bonferroni-corrected t-tests to identify individual OTUs significantly changing its relative abundance between treatments. To assess whether *L. chondrodes*’ top nine core OTUs changed their abundance ranks consistently in shaded *vs*. control samples we calculated Kendall’s rank correlation coefficient between sample pairs and tested for significant (Bonferroni-corrected) rank correlations between samples. The Sample by OTU matrix and a bash script with the *vsearch* pipeline used to generate OTUs are available in the project repository. In this repository the R scripts used to analyze the data are also available.

### Histological work

For histology, samples were fixed in a ∼4% formalin in seawater solution overnight at ∼5 °C, in 2,5% glutaraldehyde+2M sucrose in seawater or using Bouin’s solution + 2M sucrose. The tissue was dehydrated in a graded ethanol series and embedded in paraffin. Sections (5 to 15 *µ*m thick) were made and stained using DAPI/Phalloiding or Masson-Trichrom. Documentation of the sections was performed on a BH-2 light microscope (Olympus) in combination with an Axio Cam I (Carl Zeiss) or a Leica Thunder Imager.

For scanning and transmission electron microscopy samples were fixed in a fixation cocktail containing 2% glutaraldehyde and 1% osmium tetroxide in sterile filtered seawater and subsequently dehydrated in a graded ethanol series. Samples for the SEM were embedded in methacrylate. The final mixture was prepared according to (Reimer, 1967) and (Stockem & Komnick, 1970). From the hardened blocks semi-thin sections were cut with a HM 360 Mikrotom (Microm-International GmbH). These cut samples were disembedded using xylene and transferred into ethanol or acetone. The dehydrated samples were dried on an Emitech K850 Critical Point Dryer (Quorum Technologies) and gold coated on an Emitech K500 Sputter Coater (Quorum Technologies). Imaging was done using an ESEM XL30 scanning electron microscope (Philips). For ultrastructural studies, samples were embedded in Durcupan ACM (Sigma-Aldrich). Ultra-thin sections of about 50 – 70 nm were placed on Formvar coated copper grids and contrasted with uranyl acetate and lead citrate according to Reynolds (1963). Imaging was done on an EM 900 and CEM 902a transmission electron microscope (Carl Zeiss) with an accelerating voltage of 80 keV.

### Transcriptome sequencing, assembly and annotation

RNA was extracted from flash frozen sponge explants using a standard CTAB protocol followed by an isolation/clean-up using ZR-Duet™ DNA/RNA MiniPrep (Zymo). The concentration and quality of the RNA extracts was controlled on a Nanodrop (to obtain 260/280 and 260/230 ratios) and a Bioanalyzer 2100 and the extractions were used to produced libraries for RNA-Seq at the EMBL Sequencing Core. These libraries were 50bp PE sequenced, quality controlled with FastQC and used for *de novo* transcriptome assembly in Trinity version 2.0.6 (Grabherr et al., 2011). Transcriptome completeness was assessed using BUSCO (Simão, Waterhouse, Ioannidis, Kriventseva, & Zdobnov, 2015).The assembly was annotated against Swiss-Prot, the *Amphimedon queenslandica* AQU2 protein set (http://amphimedon.qcloud.qcif.edu.au/; (Fernandez-Valverde, Calcino, & Degnan, 2015) and the KEGG database (http://www.genome.jp/kegg/). For transcripts that could be annotated using the Uniprot Swiss-Prot database, the GO terms of the best Uniprot match were fetched and used. Protein domains were searched using PfamScan.pl (ftp://ftp.ebi.ac.uk/pub/databases/Pfam/Tools/). Transcripts were translated using the program Transdecoder version 2.0.1 (https://github.com/TransDecoder/TransDecoder/wiki). All annotations were compiled on a csv metatable. The assembly, the annotation files, the metatable and several scripts used to conduct the analyses are available in the project repository.

### Differential gene expression analyses and GO-term enrichment analyses

Upon assembly of the *L. chondrodes*’ reference transcriptome, the program RSEM version 1.2.28 (Li & Dewey, 2011) was used to obtain counts per transcript for each library. The count matrix generated was analyzed with the R package wgcna (Langfelder & Horvath, 2008) to determine clusters of transcripts with similar expression patterns (hereafter, transcription modules). We allowed for clusters with a minimum size of 30 transcripts and merged modules using a module eigengene expression dissimilarity threshold of 0.35. For the resulting modules, significant differences in eigengene expression between control and shaded sponges were assessed using a permutation-based Student’s t tests. For significant expression modules, we conducted GO-term enrichment analyses with TopGO (Alexa & Rahnenfuhrer, 2016) using a Fisher test on nodes with at least five annotations. To summarize the resulting list of significantly enriched GO-terms, we used the R package GOSemSim (Yu et al., 2010) to calculated the pairwise semantic similarity (Wang) between GO-terms and clustered (Ward) the enriched GO-terms into groups of semantically related GO-terms. To assign a representative GO-term to each cluster of semantically related, significantly enriched GO-terms in the different modules, we took advantage of the hierarchical structure of the gene ontology and retrieved subgraphs including the GO-terms in each cluster. These subgraphs show how semantically similar GO-terms relate to each other in the broader gene ontology directed acyclic graph. We used these subgraphs graphs to visualize significance and information content (IC) of different enriched GO-terms as well as their relationship with other enriched GO-terms. Highly significant GO-terms in a cluster were used as cluster descriptors. In addition, DESeq2 (Love, Huber, & Anders, 2014) was used to identify specific transcripts that were differentially expressed (|LFC| ≥ 1, FDR < 0.01) in control *vs*. shaded sponges and corroborate the modulation of specific pathways in shaded *vs*. control sponges. R scripts to conduct these analyses are provided in the project’s repository.

### Read archiving and project repository

All short reads generated for this study were deposited in the European Nucleotide Archive (ENA) under Bioproject PRJEB24503. In addition, the assembled and annotated reference transcriptome, and all scripts and data matrices used in this study can be retrieved at https://gitlab.lrz.de/cbas/cbas_resources

## Results

### *Shading results in the collapse of the cyanobacterial population and the reorganization of the microbiome of* L. chondrodes

Shading cyanosponges in the field reduces the abundance of cyanobacterial symbionts and affects the sponge growth rate (Thacker, 2005; Usher, 2008). Similarly, shading *L. chondrodes* causes a progressive loss of its characteristic blue/purple color and a drop in *chlorophyll-a* activity, as measured by PAM fluorometry (Curdt, Schupp, & Rohde, 2022). *Chlorophyll-a* activity drops below the PAM detection limit after 56 days of shading (Fig. 2A). High-throughput 16S V4 rRNA amplicon sequencing of foliose (i.e., blue/purple) and thread-like (i.e., white/bleached) *L. chondrodes* samples and of sponge explants exposed to zero (i.e., control explants), three, six, nine, and twelve (i.e., bleached explants) weeks of shading revealed a significant shift of the microbial community of control (foliose, blue/purple sponges, Fig. 2C) *vs*. shaded/bleached (thread-like, Fig. 2D,E) sponge explants upon exposure to shading (ANOSIM R_1000_ = 0.4234, p=0.0009). The change in the microbiome was progressive, with sponges exposed to three weeks of shading showing microbiomes more similar to those of control samples and sponges exposed to nine weeks of shading closer to those observed in bleached sponges (Fig. 2B); sponge explants exposed to six weeks of shading had microbiomes intermediate between control and bleached sponges in NMDS space. Also, *L. chondrodes* explants exposed to scattered light with photosynthetic active radiation (PAR) below detection levels showed similar microbiome changes (ANOSIM R_1000_ = −0.1676, p=0.9151) to those observed in shaded specimens but never lost their purple coloration (Fig. 2E).

**Figure 2.**
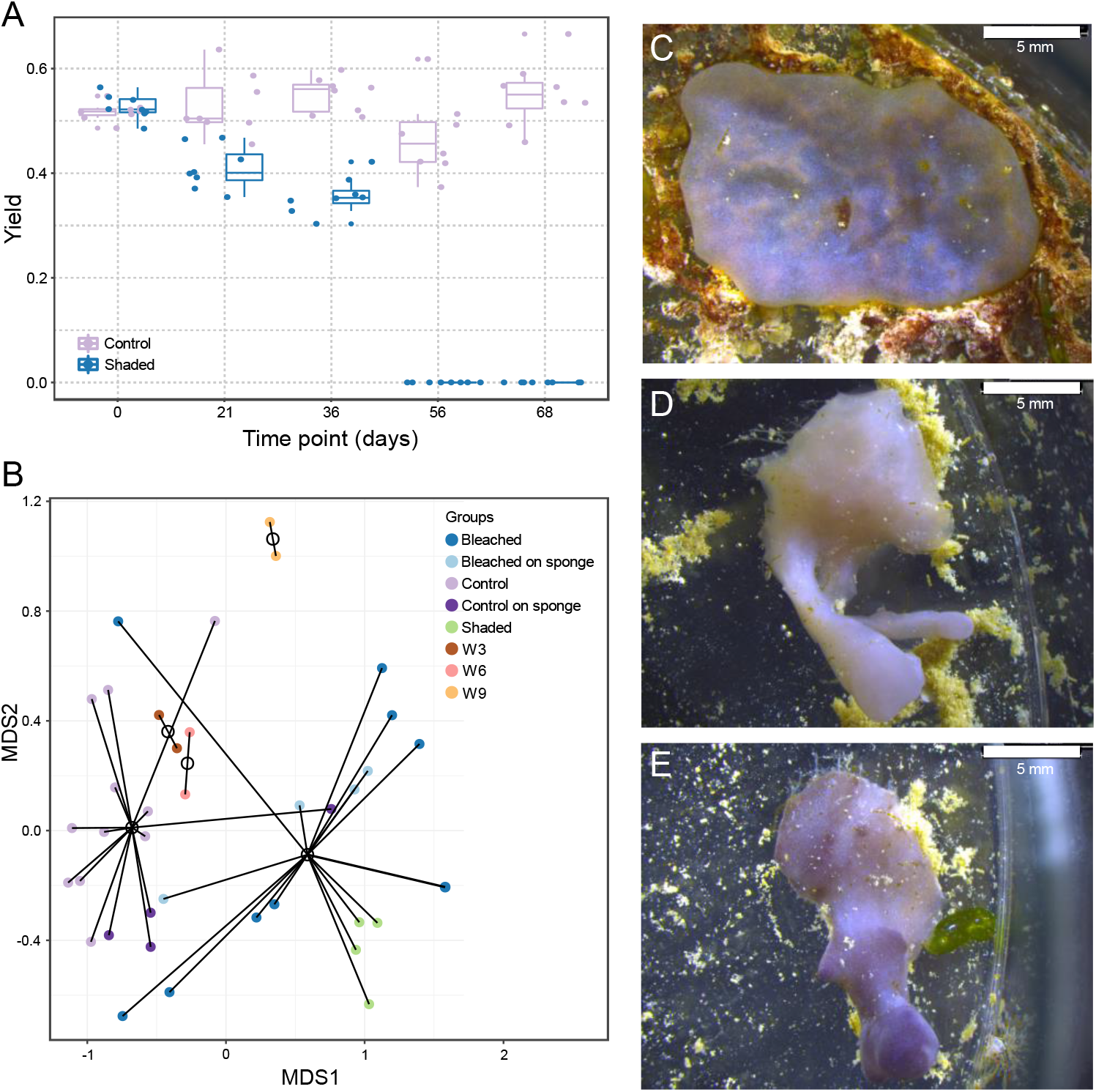
Photosynthetic yield, microbiome composition and morphological changes observed in control *vs*.shaded *L. chondrodes* explants. A. Photosynthetic yield of control and shaded sponges at different timepoints during a characteristic shading experiment. B. NMDS (Bray-Curtis dissimilarity) of control, bleached and shaded *L. chondrodes* explants as well as fragments (control and bleached) sampled directly from the sponge. C-E. *L. chondrodes* explants after 12 experimental weeks. Note the developed projections in shaded explants (D and E) which are not present in control explants (C).

In general, the community shift observed in shaded *L. chondrodes* explants was driven by the collapse of the cyanobacterial population. This bacterial group was significantly depleted (LFC = - 4.60; *t*_18.40_ = −10.97, Bonferroni corrected p-value < 0.001) in thread-like, shaded sponges compared to foliose, light-exposed sponges (Fig. 3, and Suppl. Fig. 2). In contrast, the abundance of other OTUs in *L. chondrodes*’ core microbiome (*sensu* Vargas et al. 2021) barely differed between foliose and thread-like sponges, with only an acidobacterium (OTU_3) and a proteobacterium (OTU_5) increasing their abundance (LFC = 1.95 and 0.69, respectively; *t*_17.08_ = 3.18, Bonferroni corrected p-value = 0.049 and *t*_29.81_ = 3.82, Bonferroni corrected p-value = 0.006, Fig. 3 and Suppl. Fig. 2).

**Figure 3.**
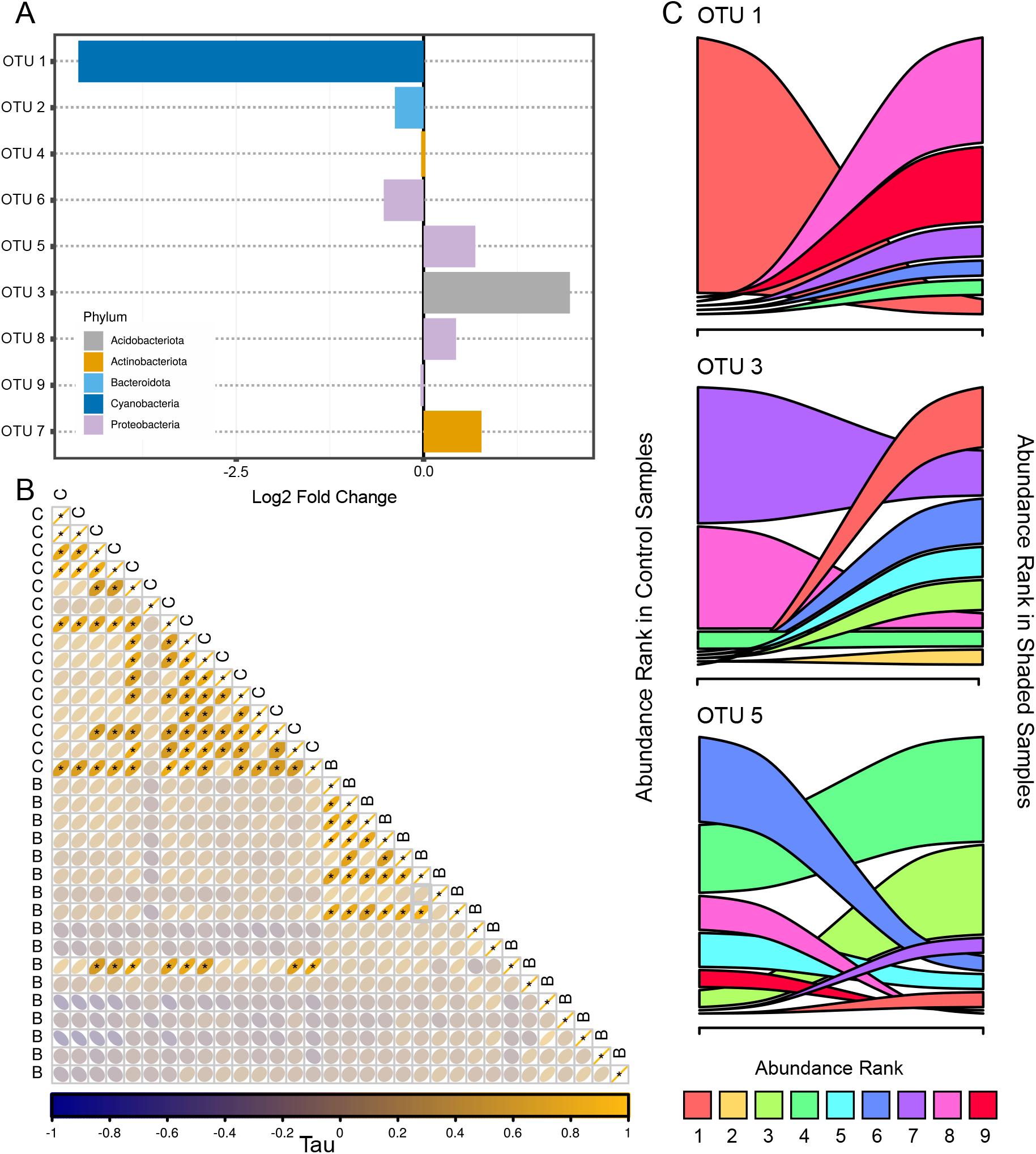
Microbiome reaccomodation during shading in *L. chondrodes*. A. Abundance Log_2_ Fold Change (LFC) for the top nine core OTU in control and shaded *L. chondrodes* sponges. B. Abundance rank correlogram of *L. chondrodes* top nine core OTUs. Note the positive rank correlations among control samples. C. Change in abundance rank for three OTUs showing a significant change in abundance between control and shaded samples. Note the generalized abundance rank decay of OTU_1 (Cyanobacteria) and the less clear but consistent increase in abundance rank of OTU_3 (Acidobacteria) and OTU_5 (Protebacteria).

Notably, the community reaccommodation observed upon shading was consistent among samples as judged by the correlation in the abundance ranks of the core OTUs measured in control and shaded samples (Fig. 3). This analysis revealed a significant correlation in the abundance ranks of the core OTUs between samples belonging either the control or the shaded group but no significant correlation in the abundance ranks of core OTUs between control and shaded samples (Fig. 3). Clear changes in rank abundance could be observed for Cyanobacteria (OTU_1), which consistently occupied the first abundance rank in control samples but dropped to the eighth and ninth abundance rank in shaded sponges (Fig. 3). Acidobacteria (OTU_3) and one proteobacterium (OTU_5) also had marked rank abundance changes. Acidobacteria frequently occupied the seventh and eighth abundance ranks in control sponges whereas in shaded samples it occupied upper abundance rank levels (*i.e*., ranks one to eight; Fig. 3). The change in abundance rank of the proteobacterium (OTU_5) in shaded samples was less clear. This OTU occupied a broader range of abundance ranks in control and shaded samples, however in shaded sponges it occupied more frequently upper abundance ranks (*i.e*., rank three; Fig. 3). All remaining OTUs either preserved their abundance rank or moved only one abundance rank in shaded compared to control samples.

### *Shading triggers a body-plan reorganization in* L. chondrodes

Shading can impair sponge growth in the field (Thacker, 2005; Usher, 2008). In *L. chondrodes*, shaded explants only increased their area by ∼15% whereas the area of control sponges increased ∼150% after 67 experimental days (Curdt et al., 2022; Fig. 4). A transition from a foliose to a thread-like morphology consistently accompanies growth stagnation and bleaching in shaded sponges (see Fig. 2). Importantly, sponges exposed to scattered light with PAR below detection levels also transition to a thread-like morphology while retaining their color (see Fig. 2). To study the effect of shading on sponge morphology we investigated the microanatomy of control and bleached sponges. Histological sections of control and shaded sponges revealed a contrasting micromorfology in these two treatment groups. Control sponges are highly organized, with a clearly delimited choanosome and a palisade of vacuole-rich cells (hereafter polyvacuolar gland-like cells, PGCs) underneath the pinacoderm (Fig. 4B, Suppl. Fig. 3). Under control conditions, PGCs occur interspersed in the pinacoderm and form pore openings between exopinacocytes (Fig. 4C-D). Notably, PGCs physically interact with and ingest bacteria in the mesohyle (Fig. 4E), where three PGC morphotypes with varying vesicle sizes (Fig. 4F-G) coexist. In contrast, shaded sponges had a disorganized micromorphology with no identifiable choanosome and a depletion of PGCs as judged by the reduction of *L. chondrodes’* characteristic subpinacodermal cell palisade (Fig 4B, Suppl. Fig. 3).

**Figure 4.**
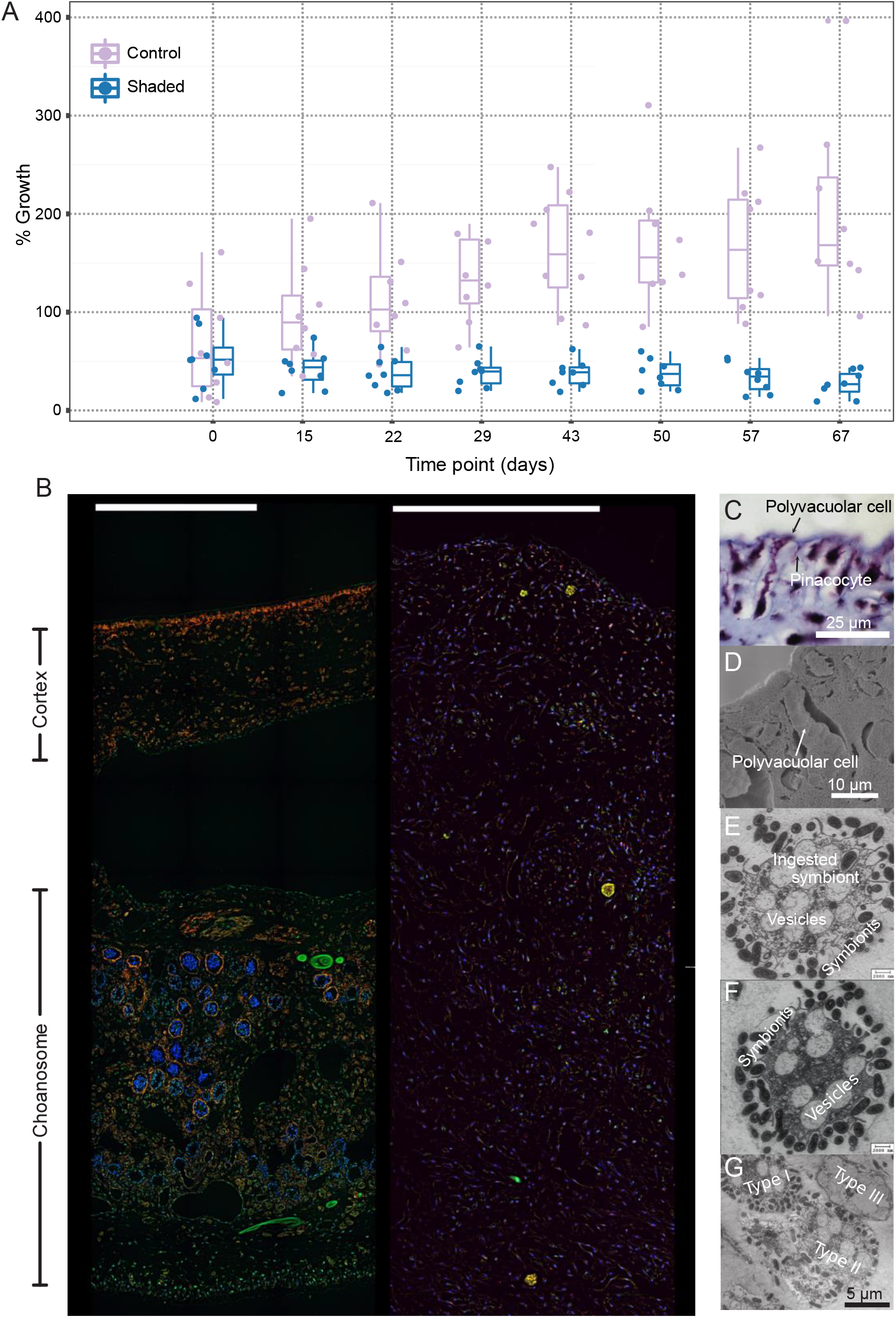
Growth and general microanatomy of control and shaded *L. chondrodes* specimens. A. Growth of control and shaded *L. chondrodes* explants. B. General microanatomy of control (left) and shaded, thread like *L. chondrodes*. Note the highly organized cortex/choanosome of control specimens *vs*. the unstructured appearance, mostly composed of dispersed cells without a clear choanosome, of shaded explants. C. Light and scanning electron microscope images of *L. chondrodes* palisade of polyvacuolar gland-like cells and transmission electron microcopy images of these cells interacting with symbiont in the sponge mesohyle.

### L. chondrodes *regulates immune and developmental coexpression modules during the foliose to thread-like morphology transition*

To evaluate whether the morphogenetic changes observed in shaded *L. chondrodes* are accompanied by a modification of the sponges’ transcriptional state, we used RNA-Seq to *de novo* assemble a reference transcriptome for *L. chondrodes*. The assembly consisted of 128,686 transcripts >200bp (N50 = 1,281bp, mean length = 803bp) and is highly complete (Suppl. Table 1). Upon translation, 37.75% of all the transcripts resulted in peptides >100 amino acids, of which 29.44% had a SwissProt blast match, and 36.21% matched a protein present in the marine sponge *Amphimedon queenslandica* (Fernandez-Valverde et al., 2015).

A comparison of the global expression patterns of shaded *vs*. control samples revealed significant changes (Adonis Pseudo-F_1,7_ =5.1164, p=0.04) in the transcriptional state of the shaded sponges. A gene coexpression network analysis resulted in 10 meta-modules with 103 to 5281 transcripts (Suppl. Fig. 4). Four meta-modules had significantly different eigengene expression values in control *vs*. shaded sponges (Suppl. Fig. 5). These meta-modules contain ∼72% (12,943) of all the transcripts analyzed (17,893) and include the three most populated meta-modules found with 5,281, 4,595, and 3,114 transcripts each. The largest significant meta-module (“lavenderblush2”) includes transcripts showing higher eigengene expression values in shaded compared to control sponges. This metamodule was enriched in transcripts involved in diverse developmental (*e.g*., the canonical Wnt pathway), signaling (*e.g*., TLR signaling pathway), cell adhesion, cell organization, cell activation, and cell differentiation processes, the regulation of apoptotic and necrotic cell death, of transcription, of protein phosphorylation and ubiquitination, and of the cells response to (inflammatory) stress and oxygen reactive species, among other (Fig. 5 and Suppl. Table 2). The remaining significant metamodules had a higher eigengene expression in control sponges. These metamodules were enriched in the regulation of metabolic and biosynthetic processes, the regulation of phosphorylation, DNA replication and repair, transcription and gene expression, cell catabolism, cell communication and mobility, development and morphogenesis, signaling and signal transduction, immune processes, the regulation of cell division and proliferation, secretion and intracellular transport and localization, and ion homeostasis (Fig. 6, Suppl. Figs 6 and 7, ans Suppl. Tables 3-5).

**Figure 5.**
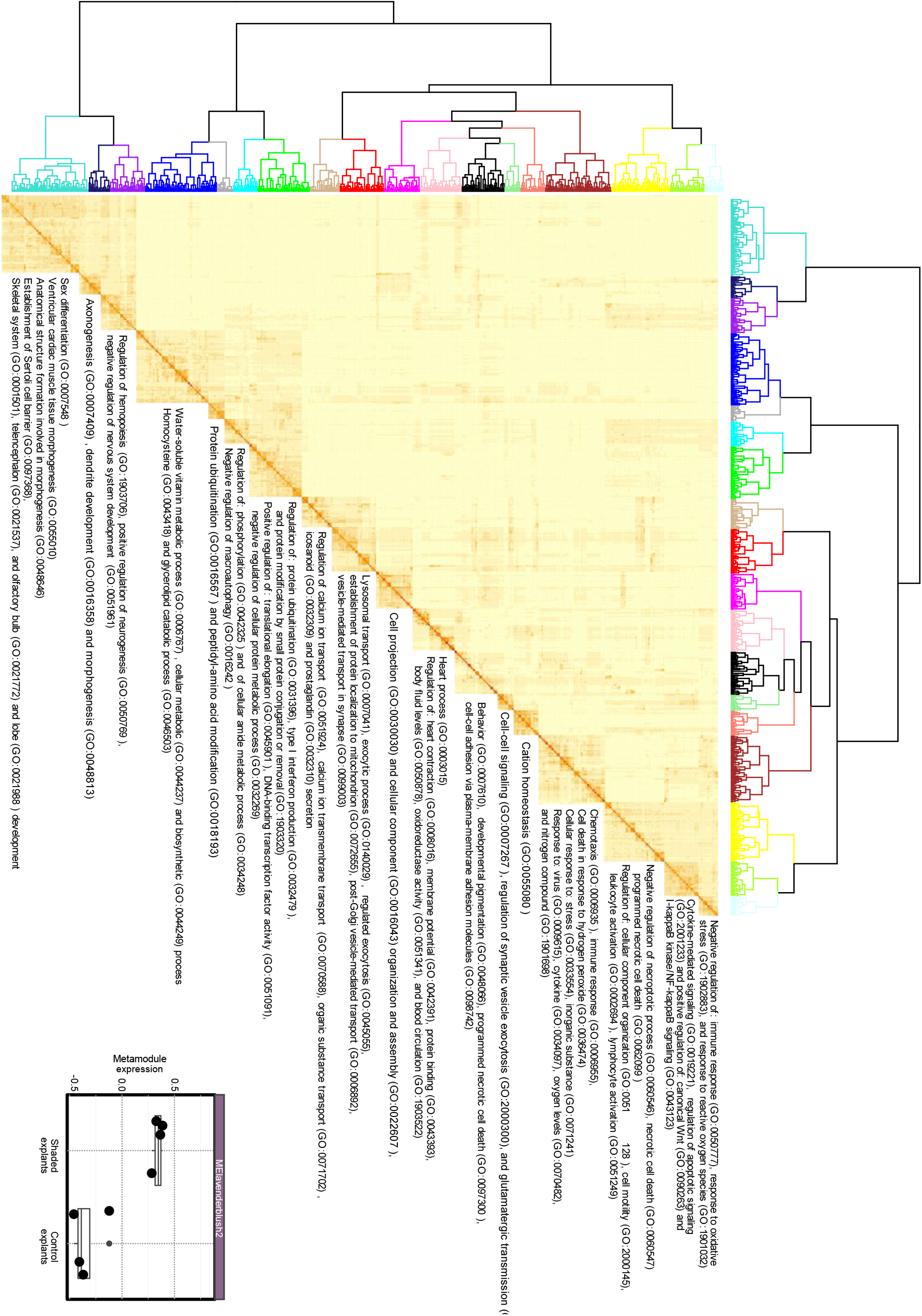
Clusters of semantically similar, significantly enriched Biological Process GO-terms in the “lavenderblush2” module and metamodule expression in control *vs*. shaded sponge explants. The representative GO-terms are listed.

**Figure 6.**
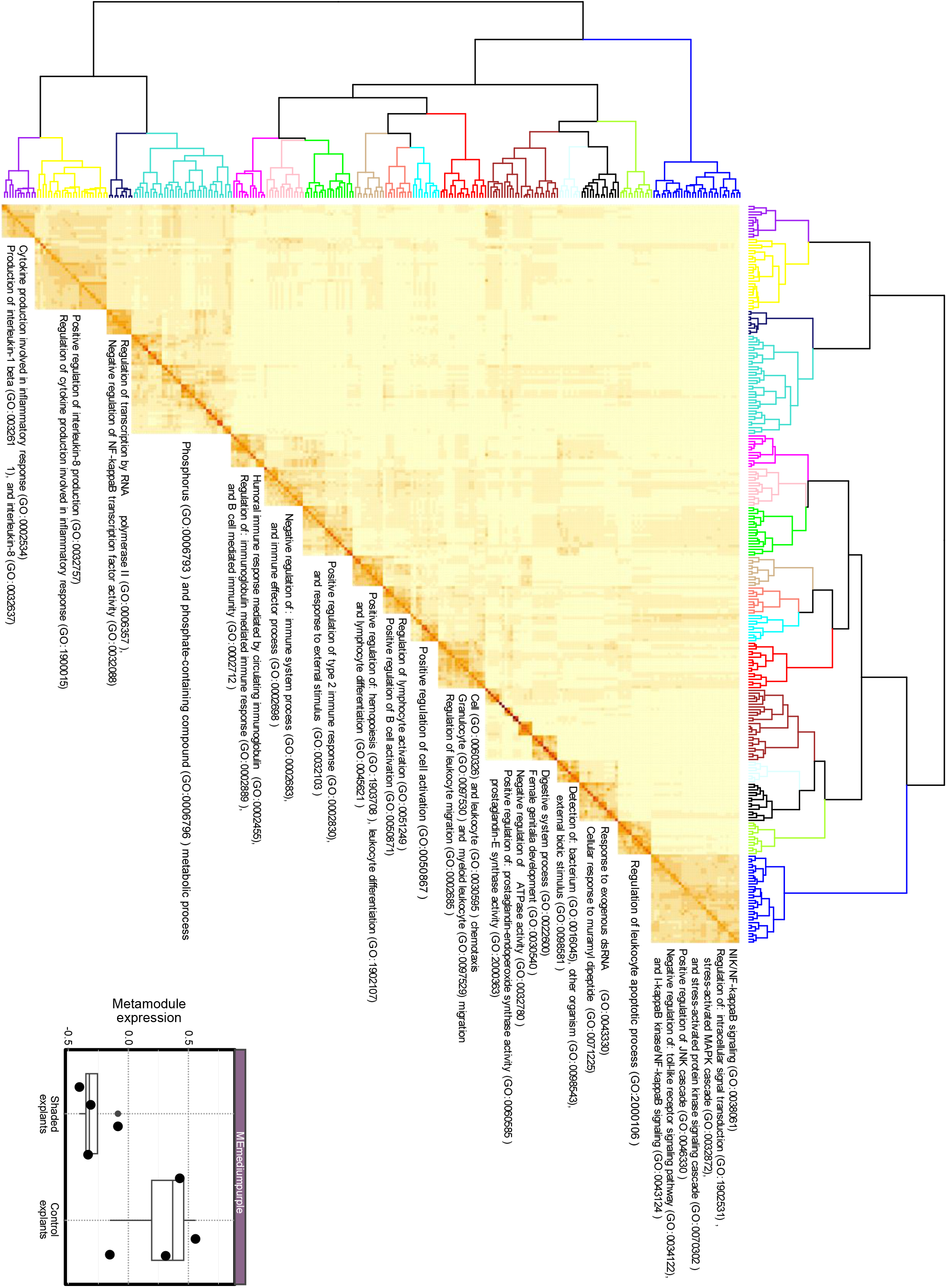
Clusters of semantically similar, significantly enriched Biological Process GO-terms in the “mediumpurple” module and metamodule expression in control *vs*. shaded sponge explants. The representative GO-terms are listed.

### Specific developmental and immune pathways are differentially modulated in foliose vs. thread-like, shaded sponges

Between treatments, a total of 5,508 transcripts were differentially expressed (BH-corrected p < 0.01). Of these, 1166 transcripts were upregulated (LFC ≥ 1) in bleached sponges compared to the control, and 2563 transcripts were downregulated (LFC ≤ −1). In agreement with our gene coexpression network analysis, GO-term enrichment analyses of this set of genes also pointed to the regulation of canonical and non-canonical Wnt signaling and, generally, to processes linked to cell development and cell movement among differentially expressed genes (Suppl. Table 6). We found an ortholog of *A. queenslandica*’s WntA and a Wnt-like transcript of undetermined phylogenetic affinity (Fig. 7 and Suppl. Fig. 8) significantly underexpressed in bleached sponges. In addition to Wnt, we found a formin-bearing transcript related to *D. melanogaster*’s MWH protein (Suppl. Fig. 9), a component of the non-canonical Planar Cell Polarity (PCP) Wnt pathway, underexpressed in bleached sponges. Finally, bleached sponges also downregulated a putative Transforming Growth Factor *β* (TGF-*β*) transcript and a putative ortholog of *Amphimedon*’s TLR receptor 1 (AmqIgTIR1) and upregulated a TNF receptor-associated factor (TRAF) and an ortholog of *A. queenslandica*’s interferon regulatory factor 1 (IRF1; Fig. 7 and Suppl. Figs. 10-13).

**Figure 7.**
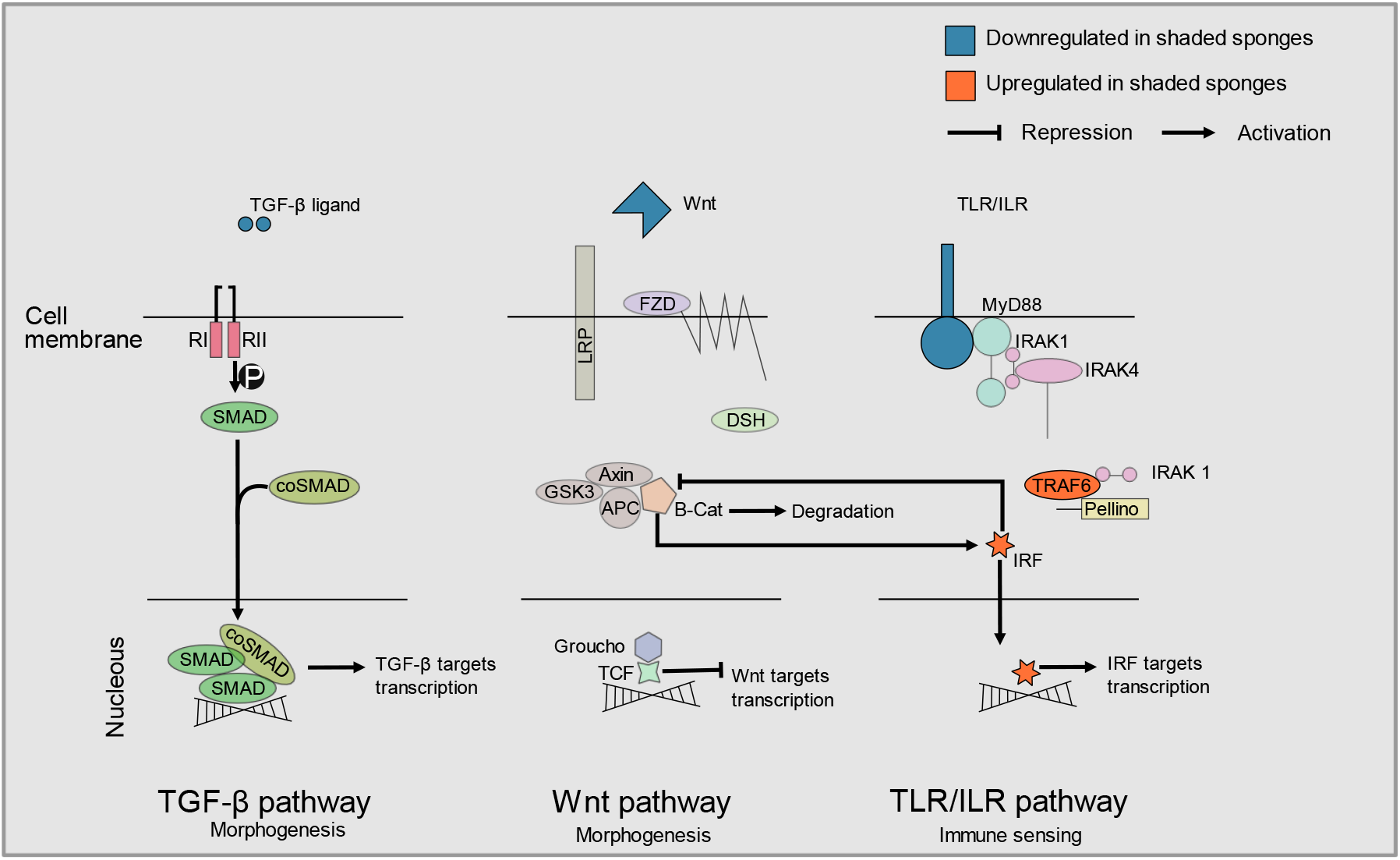
Up- and downregulated components of selected morphogenetic and immune pathways identified in control *vs*. shaded, thread-like *L. chondrodes*. These pathways were chosen based on published studies showing their importance during development in sponges.

## Discussion

Broad, coupled morphological and transcriptomic changes correlated with the loss of a key symbiont and the concomitant microbiome change have never been reported in sponges. In other animals, changes in the microbiome correlate with behavioral and developmental alterations (Cheesman et al., 2011; Klimovich et al., 2017; Sampson & Mazmanian, 2015; Sharon et al., 2016; Shin et al., 2011). Our results in *L. chondrodes* provide evidence for a correlated response of the sponge host and its microbiome to environmental perturbations. The stark foliose to thread-like transition of the sponge body-plan occurs in parallel to the changes in the microbiome caused by light deprivation and entails the modulation of diverse developmental, signaling, behavioral, and metabolic processes. It also correlates with the depletion of PCGs in bleached *vs*. control sponges. The reduction of the PCG population is of particular interest as these cells physically interact in the mesohyle with the microbiome and can, therefore, mediate host-symbiont interactions in *L. chondrodes*.

It is important to note that light deprivation, a possible confounding factor in our experimental setting, cannot explain the observed response of the host as sponges exposed to scattered light also transition to a thread-like morphology while retaining their color. As in bleached specimens, the loss of symbiotic cyanobacteria and the readjustment of the abundances of other non-photosynthetic bacterial symbionts drove the microbiome change observed in these sponges, which also consistently transition to a thread-like morphology during the treatment. Maldonado & Young (1998) observed a similar reaction of the sponge host in keratose sponges transplanted from shallow to deeper waters with low light availability. Notably, those authors only observed changes in sponge morphology in cyanobacteria-depleted specimens, in agreement with our results. These observations and the results of our manipulative experiments provide strong evidence for an active reaction of sponges that correlates with the decay in photosymbiont abundance.

Although the precise causal mechanism coupling microbiome changes with the observed reaction by the sponge host remains to be determined, our results suggest that both direct “immune” and indirect “sensing”, for instance through the perturbation in the sponge’s carbon budget linked to a smaller, photosynthetically compromised cyanobacterial population, are active in *L. chondrodes*. In this regard, we observed an active regulation of many metabolism-related GO-terms in *L. chondrodes*. Coupled with the observation that shading causes growth to stagnate in shaded sponges, in agreement with previous field experiments on different cyanosponges (Thacker, 2005), the observed active change in metabolic regulation in bleached *vs*. control sponges likely represent a compensatory reaction of the sponge to the reduction in photosymbiont-derived derived carbon upon shading (Hudspith et al., 2022; Maldonado & Young, 1998). Supporting this interpretation, we found meta-modules enriched in GO-terms involved in the sponges’ response to oxygen levels, to oxidative stress, and to reactive oxygen species as expected if sponges increased their metabolic rate upon symbiont depletion and, consequently, had a higher oxygen demand.

Direct “immune” recognition of the symbiotic bacteria, for instance by PCGs, could be in place in *L. chondrodes*, and generally in sponges (Degnan, 2015; D. Reynolds & Thomas, 2016; Yuen, Bayes, & Degnan, 2014). Immune processes have been previously found to be modulated during bleaching and reinfection in the *Symbiodinium*-associated sponge *Cliona varians* (Riesgo et al., 2014), and sponges exposed to different microbial-associated molecular patterns activate genes associated with immune responses and animal-microbe interactions (Pita, Hoeppner, Ribes, & Hentschel, 2018). In agreement with these findings, our results also revealed the modulation of transcription modules enriched in immune and developmental processes during the microbiome/morphogenetic transition caused by shading in *L. chondrodes*. In this regard, the modulation of specific elements of the Wnt, the TGF-β, the TLR, and the IRF pathways observed in *L. chondrodes* appear to be of particular interest. TGF-β is an important regulatory element of the immune response and is involved in multiple developmental processes, including the establishment of the animal body axes (Travis & Sheppard, 2014). During sponge development, the expression of the TGF-β ligand is highly dynamic (Adamska et al., 2007; Leininger et al., 2014) and in *S. ciliatum* adults TGF-β ligands are coexpressed with Wnt ligands around osculae and in other parts of the aquiferous system (Leininger et al., 2014). TLR signaling plays roles in development and innate immunity in the starlet sea anemone (Brennan et al., 2017) and the demosponge *A. queenslandica* (Gauthier, Du Pasquier, & Degnan, 2010). In *Amphimedon*, elements of the TLR signaling pathway are dynamically expressed during embryogenesis and several of its components are coexpressed in larval sensory cells (Gauthier et al., 2010). Bacterial modulation of the Wnt pathway via Myd88, an adaptor protein in the innate immune Toll-like receptor (TLR) signaling pathway, has been reported in zebrafish providing a mechanism whereby immune sensing and morphogenesis could be coupled in animals in general (Cheesman et al., 2011). Also, IRFs are transcription factors involved in the modulation of the immune response in animals and crosstalk with the Wnt signaling pathway during the differentiation of immune cells from haematopoietic stem-cells (HSCs) (Cohen et al., 2015; Scheller et al., 2013). In this context, Wnt regulates the differentiation of HSCs in a dosage-dependent manner producing different cell types as the expression of Wnt increases (Luis et al., 2011). In mouse, a regulatory circuit involving Wnt and Irf8, a target gene of β-catenin which in turn regulates the nuclear accumulation of this protein, has been described mechanistically coupling TLR and Wnt signaling during HSC differentiation (Scheller et al., 2013). Wnt ligands activate at least three signaling cascades that are involved in different morphogenetic processes in animals (Komiya & Habas, 2008). In sponges, Wnt plays a role in the development of the aquiferous system in a homoscleromorph (Lapébie et al., 2009) and a freshwater demosponge (Schenkelaars et al., 2016; Windsor & Leys, 2010), and is expressed around osculae in the calcareous sponge *Sycon ciliatum* (Leininger et al., 2014) and the demosponge *Halisarca dujardini* (Borisenko, Adamski, Ereskovsky, & Adamska, 2016). Our data suggests that similar regulatory circuits coupling, for instance, TLR and Wnt signaling may exist in sponges opening new research avenues to further comprehend how sponge cells interact with their microbiomes and how these interactions affect the transcriptional regulation of sponges.

Broad, coupled morphological and transcriptomic changes associated with the loss of a key symbiont and the concomitant microbiome change have never been reported in sponges. In other animals, changes in the microbiome have also been associated with behavioral and developmental alterations (Cheesman et al., 2011; Klimovich et al., 2017; Sampson & Mazmanian, 2015; Sharon et al., 2016; Shin et al., 2011). Our results in *L. chondrodes* provide evidence of link between the sponge body-plan and metabolism and the state of its microbiome, implying that sponges can directly or indirectly detect and actively respond to changes in it. The co-regulation of transcripts involved in pattern recognition via TLR signaling (*e.g*., AmqIgTIR1 and IRF orthologs) and in morphogenesis, through the modulation of Wnt and TGF-β signaling, offers a possible molecular mechanism whereby sensing of symbionts by the sponge innate immune system can be coupled with Wnt and TGF-β mediated cell differentiation and cell motility to actively and dynamically reorganize the sponge’s morphology and growth to avoid environmental conditions (e.g. shading) unfavorable for its microbiome. In addition, the necessary metabolic reaccommodation of the host to compensate the loss of an important carbon source (*i.e*., the cyanobacterial population) can lead to the transcriptional modulation of different cellular process to cope with, for instance, a higher oxygen demand. In conjunction, these results suggest a deep evolutionary origin of the molecular crosstalk mechanisms used by animals to interact with their microbiomes. They also suggest that universal metazoan signaling pathways (*e.g*., Wnt, IRF, TLR, and TGF-*β*) may mediate host-symbiont interactions in sponges as it has been already proposed for other animal groups (e.g., Brennan et al., 2017; Cheesman et al., 2011; Daniel, Ball, Besselsen, Doetschman, & Hurwitz, 2017; Franzenburg et al., 2012; Tang et al., 2017). Future studies dissecting the involvement of these pathways in the crosstalk between sponges and their microbiomes are required to assess whether the genetic toolkit necessary to support complex microbiomes was already present in the last common ancestor of animals.

## Supporting information

Vargasetal_Supplementary_Figures

Vargasetal_Supplementary_Table_01

Vargasetal_Supplementary_Table_02

Vargasetal_Supplementary_Table_03

Vargasetal_Supplementary_Table_04

Vargasetal_Supplementary_Table_05

Vargasetal_Supplementary_Table_06

## Acknowledgements

We thank Gabriele Büttner for support during laboratory work, Dr. Nora Dotzler for preparing the histological sections and Dr. Peter Naumann for supporting the aquarium facilities of the Section of Geobiology and Paleobiology of the Dept. of Earth & Environmental Sciences. We would also like to thank René Neumaier for the design, administration and support of our High-Performance Computing system, our work would be impossible without his careful and detailed work. An LMU Excellent Junior Funds grant to SV partially supported this project. GW acknowledges funding through the LMU Munich’s Institutional Strategy LMUexcellent within the framework of the German Excellence Initiative. MA acknowledges funding from the Australian Research Council through Future Fellowship (FT160100068) and Centre of Excellence for Coral Reef Studies (CE140100020) grants.. SV thanks N. Villalobos T., M. Vargas V., S. Vargas V. and S. Vargas V. for their support.

## Author Contributions

**SV** Conceptualization, Methodology, Software, Validation, Formal Analysis, Investigation, Resources, Data Curation, Writing - Original Draft, Visualization, Supervision, Funding Acquisition. **LL** Investigation, Formal Analysis. **ME** Investigation, Formal Analysis, Writing - Review & Editing, Visualization. **FC** Investigation. **SR** Investigation, Supervision. **CA** Investigation. **MN** Supervision, Funding Acquisition. **PS** Supervision, Funding Acquisition. **WDO** Investigation, Writing - Review & Editing. **MA** Investigation, Writing - Review & Editing. **GW** Resources, Writing - Review & Editing, Funding Acquisition.

