## Supplementary material for "Body-plan reorganization in a sponge correlates with microbiome change": Vargasetal_Supplementary_Figures

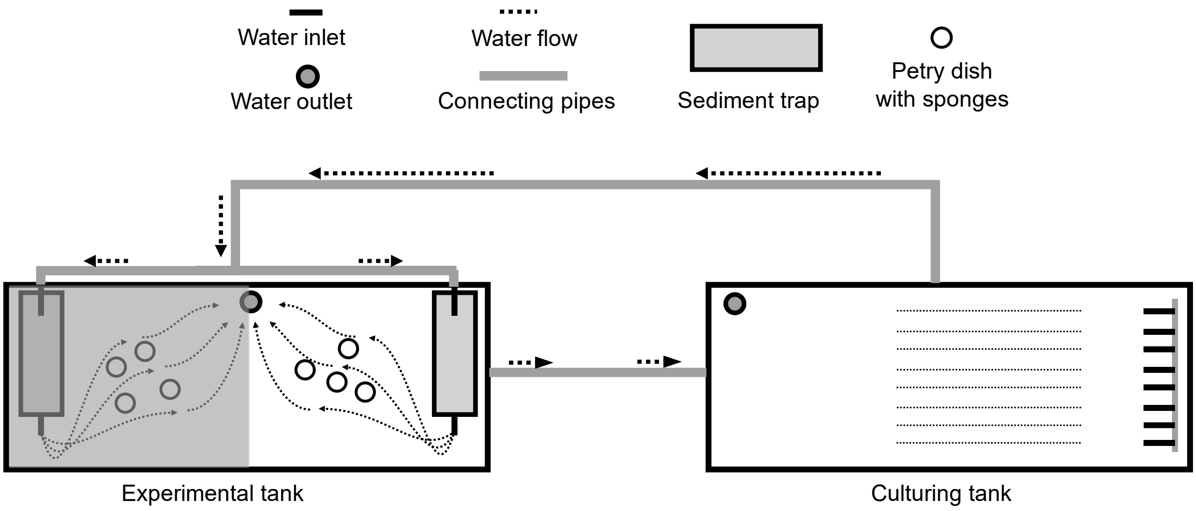


**Supplementary Figure 1.** Schematic representation of the experimental system. In the experimental tank, one half of the aquarium is covered by a black plastic cover (represented as a grey square) to shade the sponges.

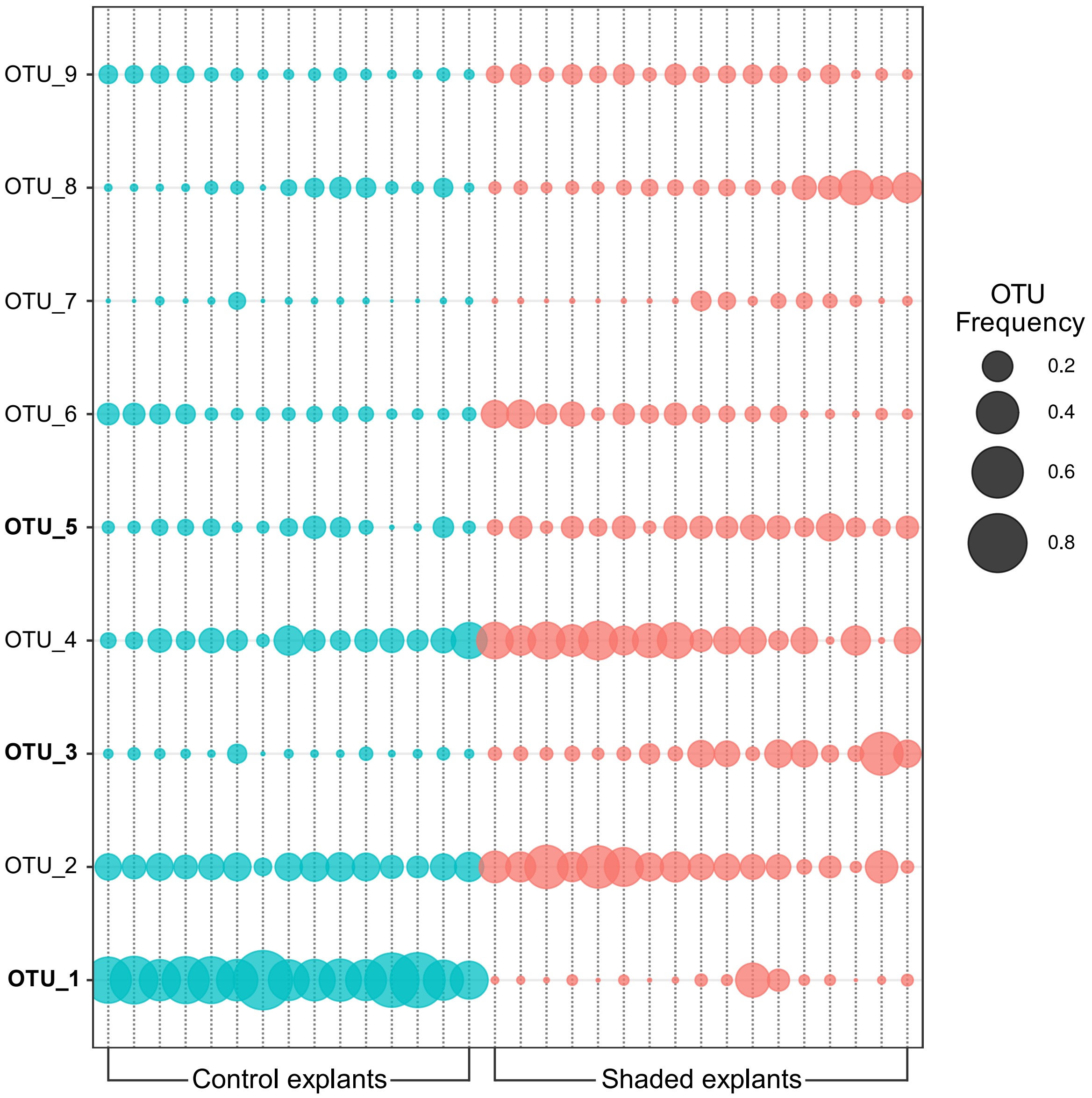


**Supplementary Figure 2**. OTU frequency by sample in control and shaded explants. OTUs significantly changing their frequency between treatments in boldface.

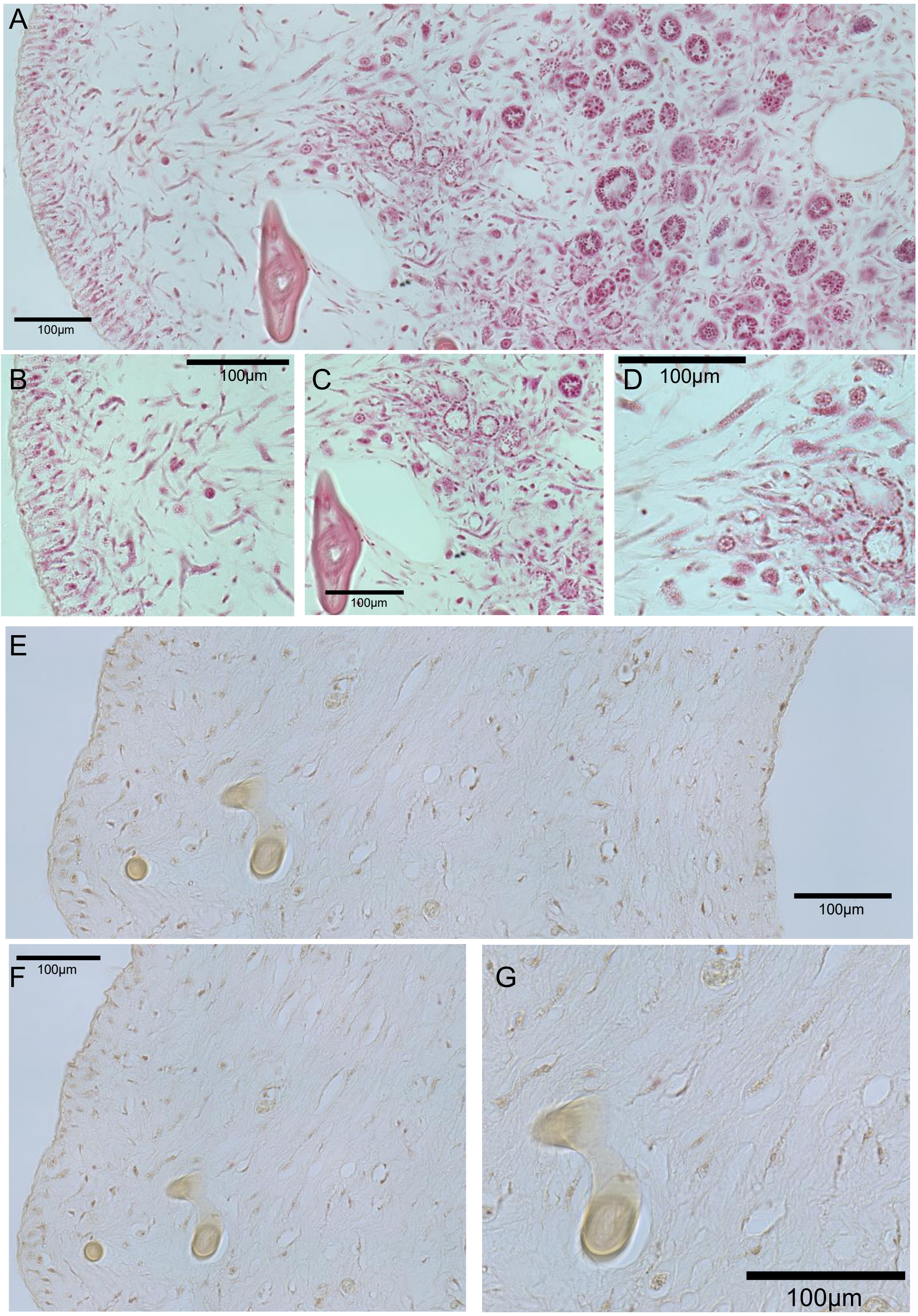


**Supplementary Figure 3**. Micromorphology of control and shaded *Lendenfeldia chondrodes*. **A**. overview of a control explant. **B**. Detail of the palisade of PGCs underneath the pinacoderm. **C**. Detail of the choanosome showing three choanocyte chambers and other cell types in the mesohyle. **D**. choanocyte chambers surrounded by different types of PGCs. **E**. overview of a shaded explant. **F**. detail showing the depleted palisade of PGCs underneath the pinacoderm. **G**. detail of the choanoderm depleted in choanocyte chambers but still showing channels and PGCs.

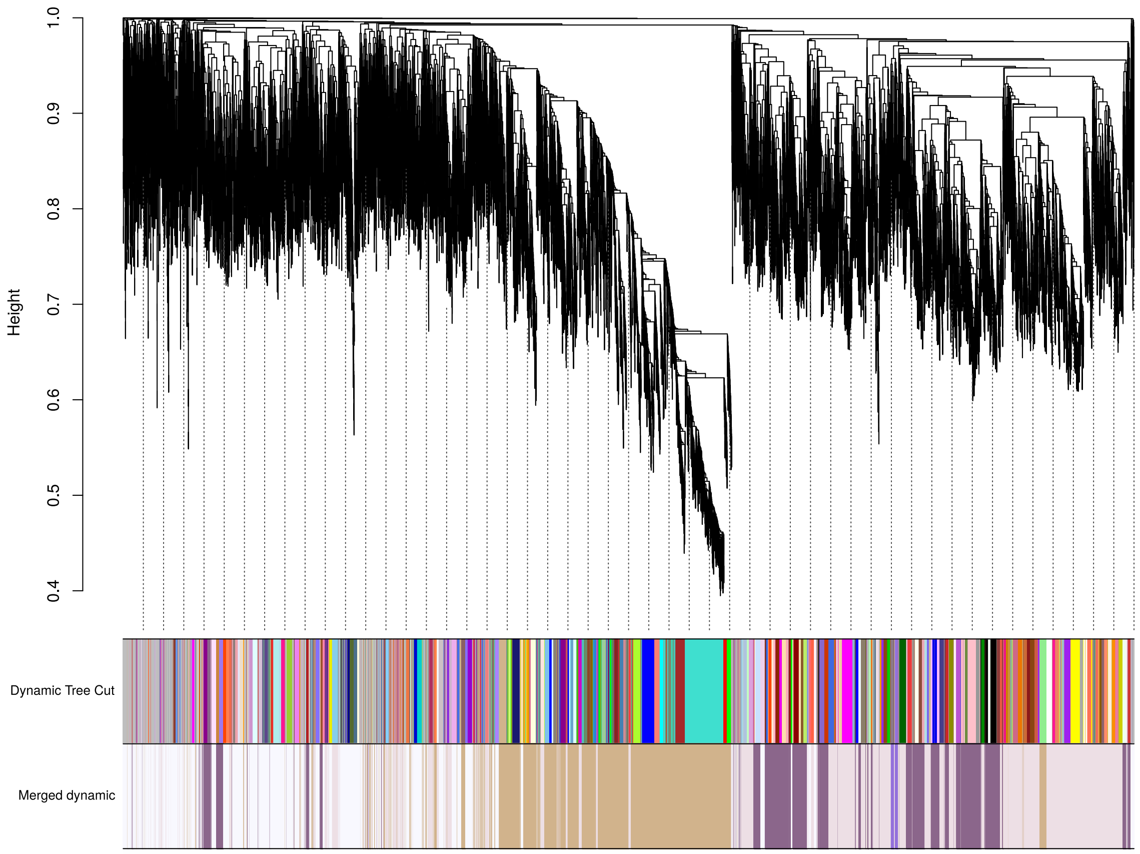


**Supplementary Figure 4**. Weighted correlated network analysis dendrogram showing the results of the dynamic tree cut + dynamic merge algorithms resulting in four main metamodules colored “tan”, “plum4”, “mediumpurple”, and “lavenderblush2”.

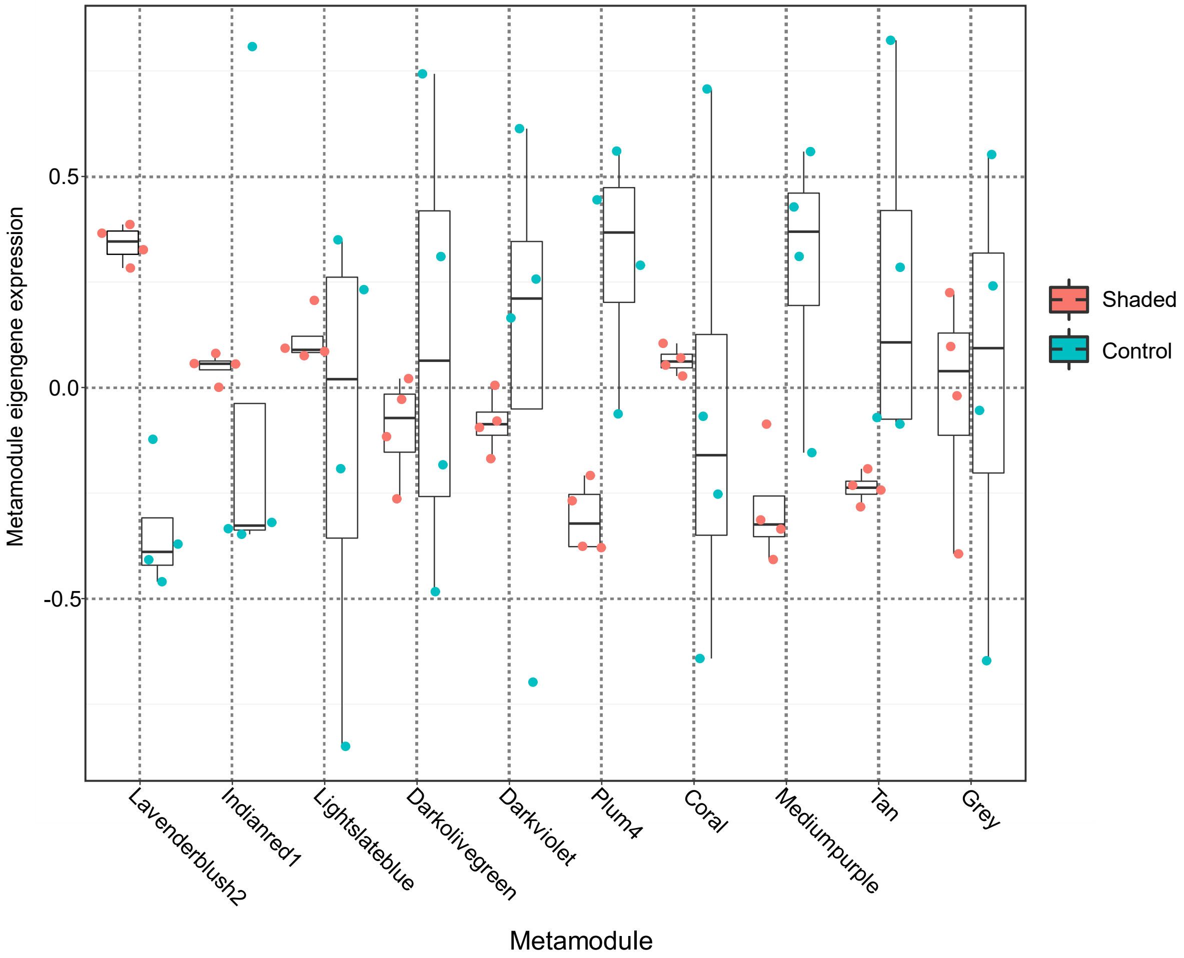


**Supplementary Figure 5**. Metamodule eigengene expression in shaded and control *Lendenfeldia chondrodes* explants. Modules showing a significantly different eigengene expression between control and treatment explants (*i.e.*, Lavenderblush2, Mediumpurple, Plum4, and Tan) are discussed in the main text.

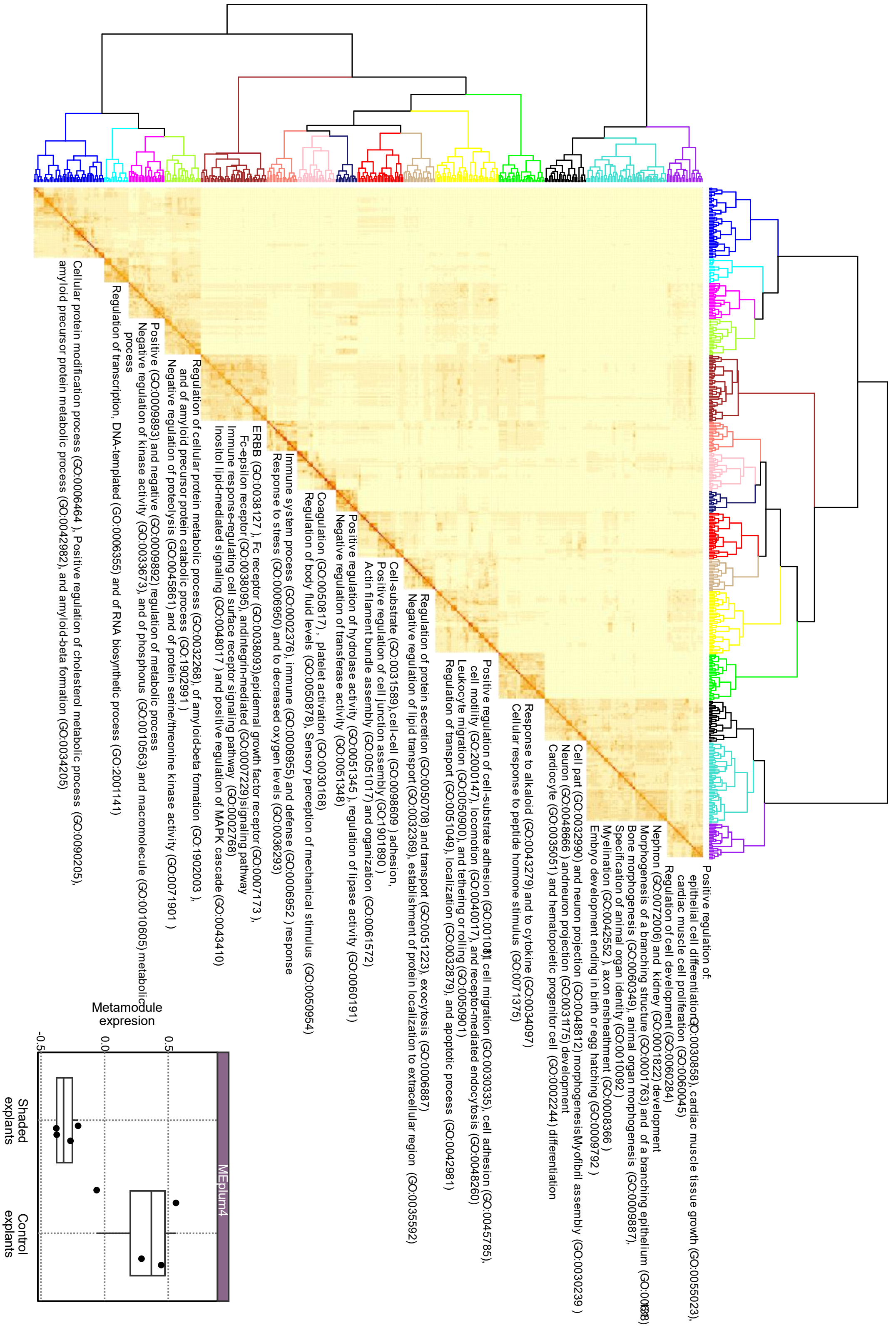


**Supplementary Figure 6**. Clusters of semantically similar, significantly enriched Biological Process GO-terms in the "plum4" module and metamodule expression in control *vs.* shaded sponge explants. The representative GO-terms are listed. See Suppl. Table X for a full list of GO-Terms in the module.

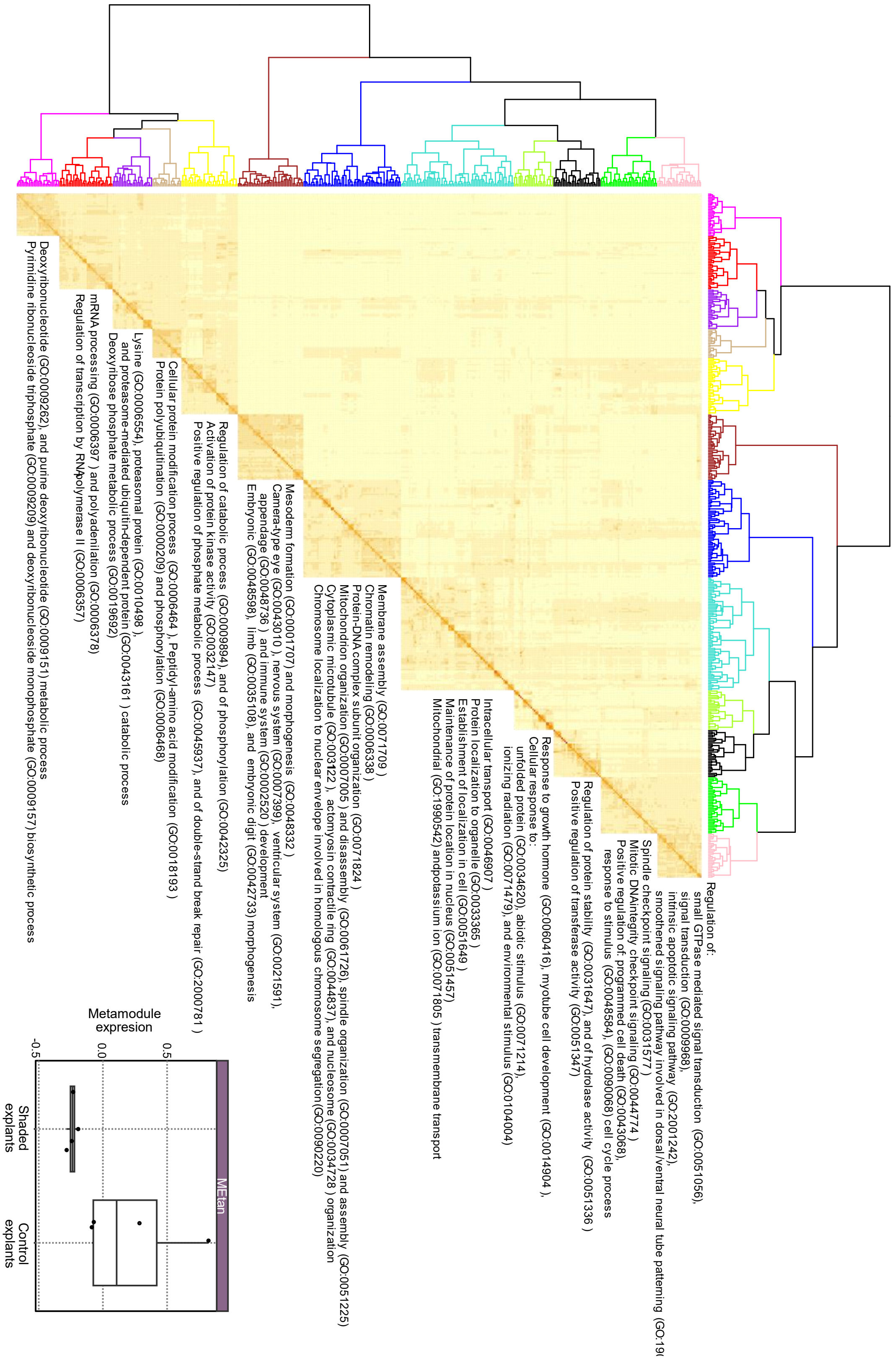

**Supplementary Figure 7**. Clusters of semantically similar, significantly enriched Biological Process GO-terms in the “tan” module and metamodule expression in control *vs.* shaded sponge explants. The representative GO-terms are listed. See Suppl. Table X for a full list of GO-Terms in the module.

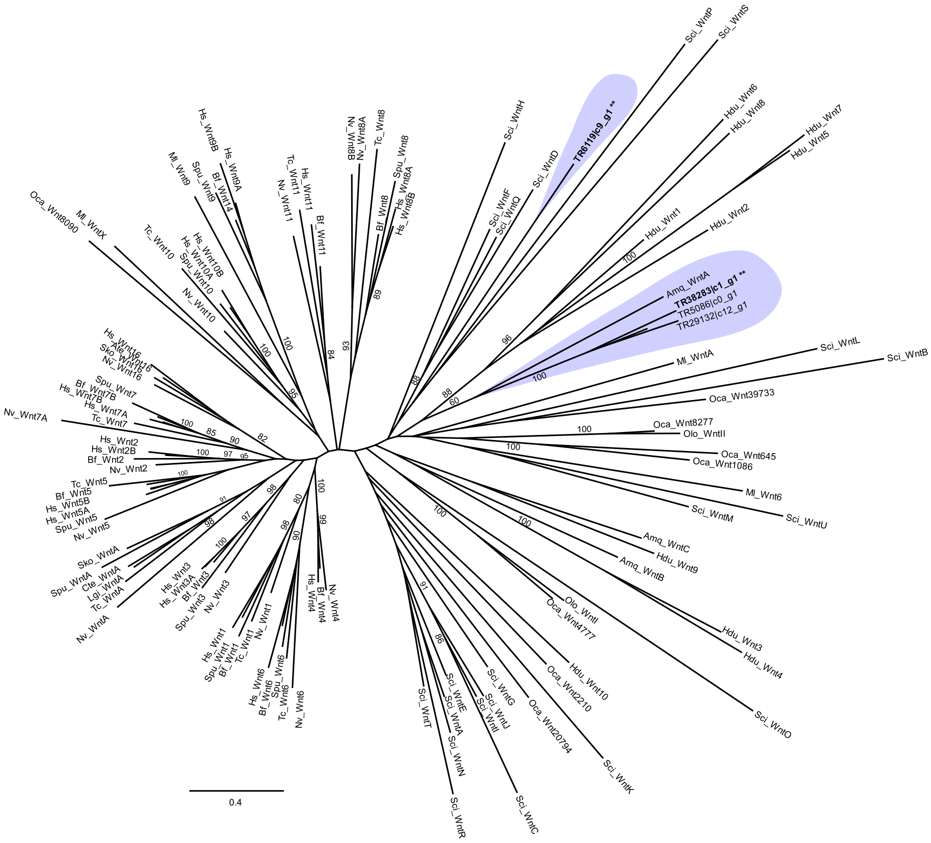


**Supplementary Figure 8**. Maximum Likelihood unrooted phylogeny of Wnt ligand proteins. The phylogeny was inferred in PhyML using the LG model. To assess support 100 bootstrap replicates were done. Two letter codes used to label different Wnt proteins as in (Borisenko et al. 2016) except for *Lendenfeldia chondrodes* transcripts, which are only referred to by their Trinity transcript name (TRXXXX) to keep the labels consistent with the provided assembly. Support values are only shown for branches with bootstrap support ≥ 70 except for the branch leading to the *A. queenslandica* + *L. chondrodes* Wnt clade. Differentially expressed (underexpressed in shaded sponges) *L. chondrodes* transcripts in bold and labeled with asterisks. The alignment used to infer this tree and a newick version of this tree is available at the project repository.

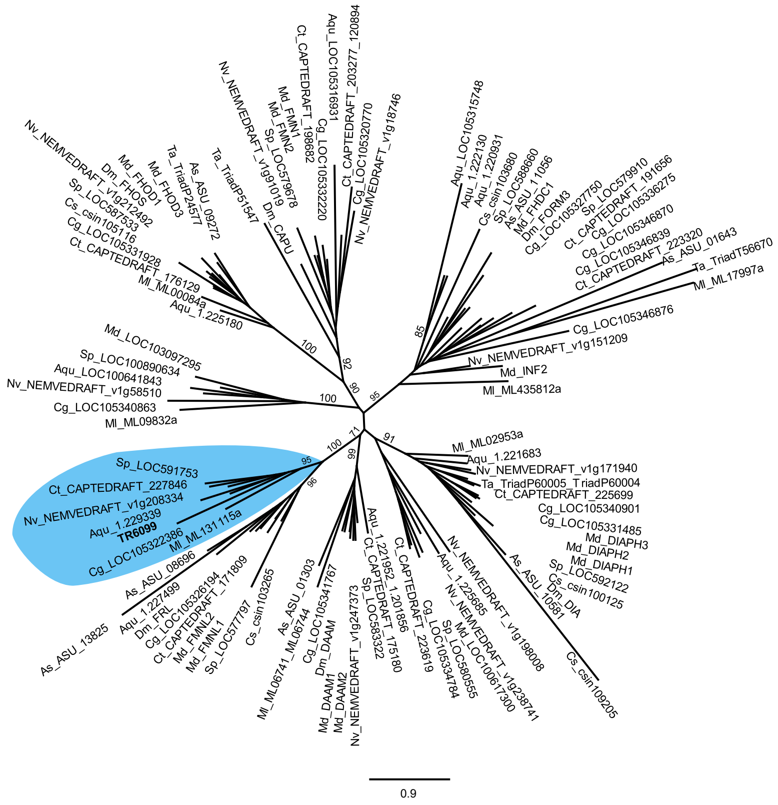


**Supplementary Figure 9**. Maximum Likelihood unrooted phylogeny of Formin bearing proteins. The phylogeny was inferred in PhyML using the LG model. To assess support 100 bootstrap replicates were done. Leaf names as in (Pruyne 2016) except for *Lendenfeldia chondrodes* transcripts, which are only referred to by their trinity transcript name (TRXXXX) to keep the labels consistent with the provided assembly. Support values are only shown for branches with bootstrap higher or equal than 70 separating major groups of proteins. The alignment used to infer this tree was modified from 26. The alignment and a newick version of this tree is available at the project repository.

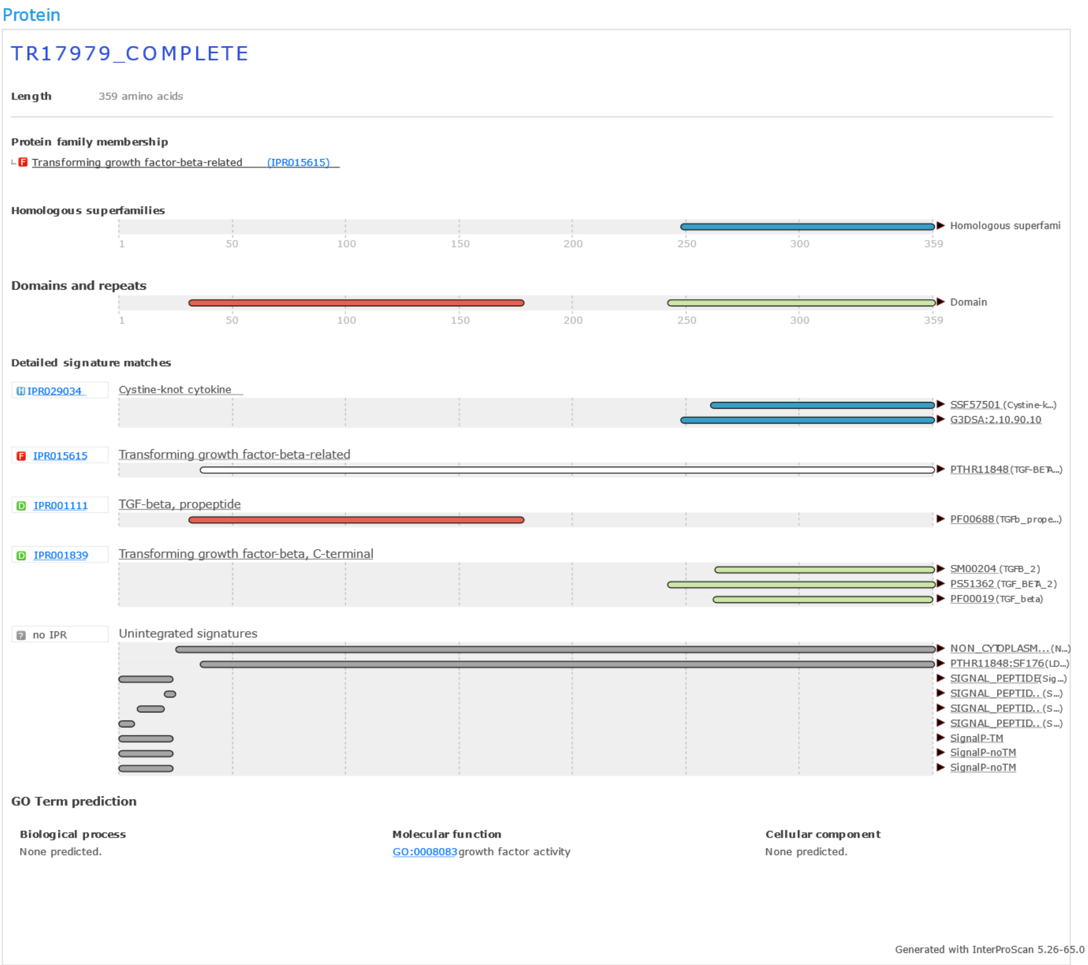


**Supplementary Figure 10**. Interproscan results for transcript TR17979. The entire peptide, not only the signaling domain, was used for the prediction. The predicted domain organization is similar to that of the human Transforming growth factor beta 1 (TGFB1) and *Amphimedon queenslandica* TGF-ß, consisting of a propeptide and a transforming growth factor-beta domain. We were unable to identify the cleavage site for this sequence. The phylogenetic position of this protein was uncertain mainly due to the fact that the Maximum likelihood phylogeny of TGF-beta proteins based only on their signaling domains is mostly unresolved (results not shown).

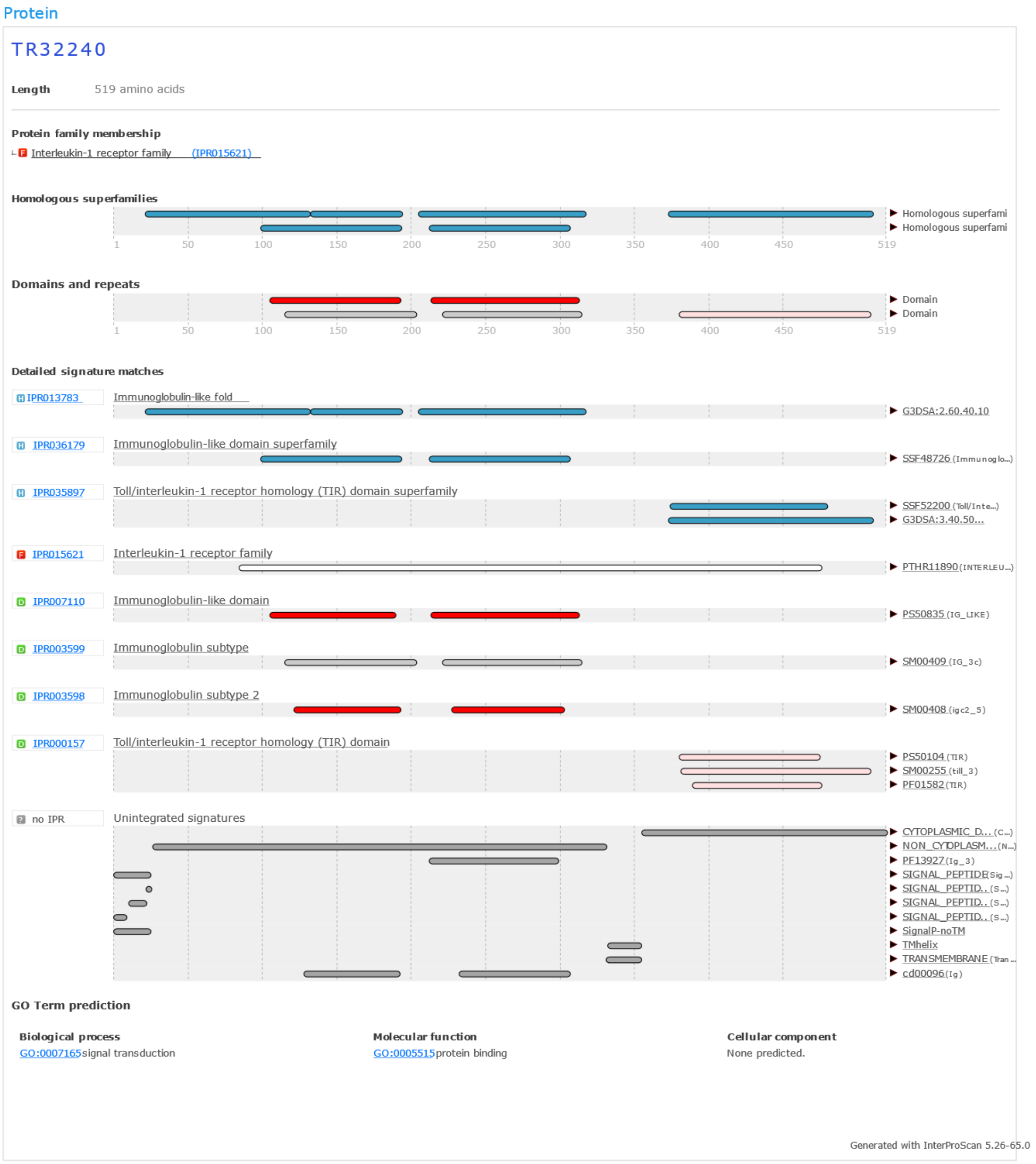


**Supplementary Figure 11**. Results of Interproscan for transcript TR32240, a putative ortholog of TLR receptor 1 in *Amphimedon queenslandica* found to be underexpressed in shaded sponges. The entire predicted peptide was scanned. The predicted domain organization, composed of N-terminal immunoglobulin-like domains and a C-terminal Toll/Interleukin-1 receptor homology (TIR) domain is identical to that of *A. queenslandica*.

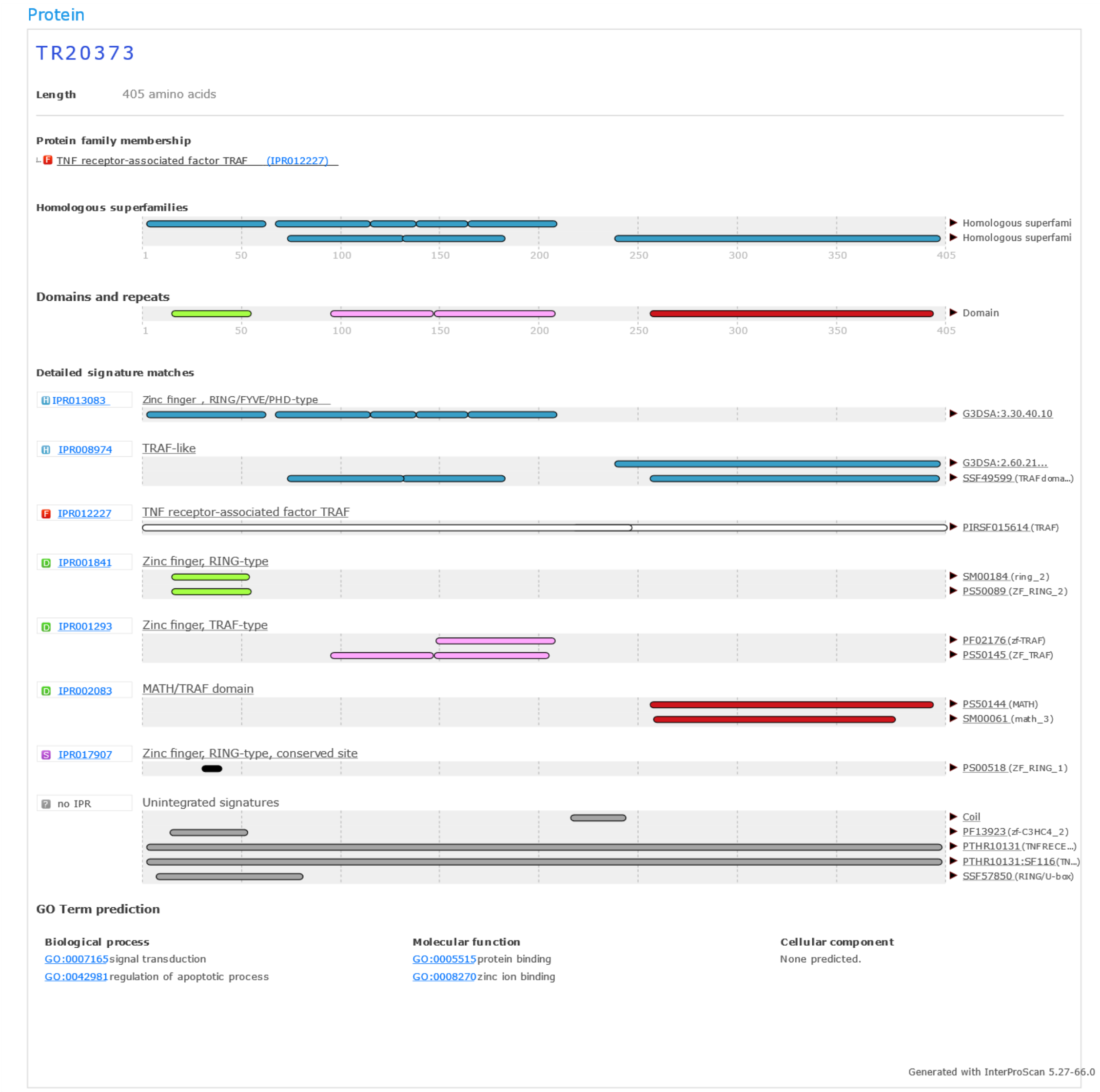


**Supplementary Figure 12**. Results of Interproscan for transcript TR20373, a putative ortholog of *Amphimedon queenslandica* TRAFs found to be overexpressed in shaded sponges. The entire predicted peptide was scanned. The predicted domain organization of this protein is similar to that of *A. queenslandica* TRAFs, consisting of N-terminal zinc finger repeats and a C-terminal MATH/TRAF domain.

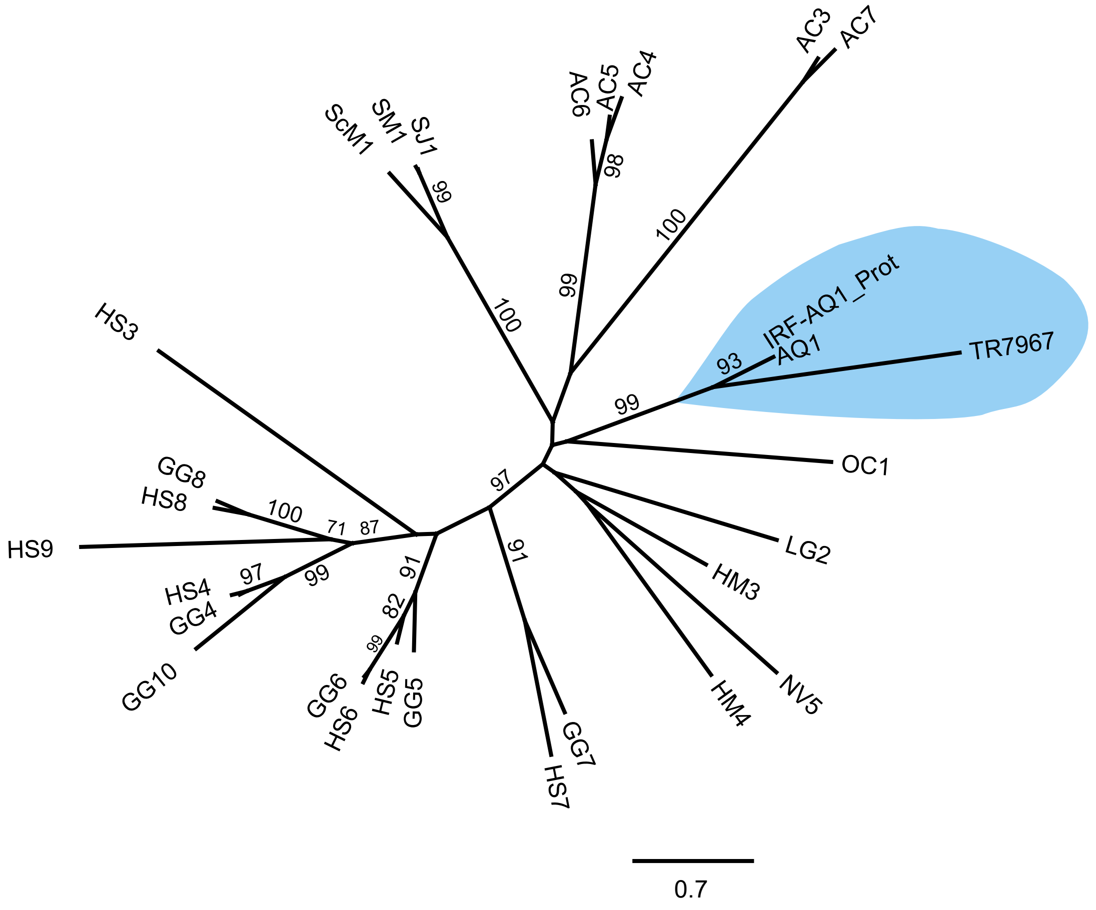


**Supplementary Figure 13**. Maximum Likelihood unrooted phylogeny of IRF transcription factors. The phylogeny was inferred in PhyML using the LG model. To assess support 100 bootstrap replicates were done. Leaf names as in (Nehyba, Hrdlicková, and Bose 2009) except for *Lendenfeldia chondrodes* transcripts, which are only referred to by their trinity transcript name (TRXXXX) to keep the labels consistent with the provided assembly. Support values are only shown for branches with bootstrap higher or equal than 70. The alignment used to infer this tree was modified from 27. The alignment and a newick version of this tree is available at the project repository.
