## Supplementary material for "Body-plan reorganization in a sponge correlates with microbiome change": Vargasetal_Supplementary_Table_01

**Supplementary Table 1**. BUSCO results for the reference transcriptome of *Lendenfeldia chondrodes*.

| BUSCO category | Total found | Percentage |
|---|---|---|
| Complete and Single Copy | 579 | 59.2% |
| Complete and Duplicated | 316 | 32.3% |
| Fragmented | 38 | 3.9% |
| Missing | 45 | 4.6% |

Total BUSCOs = 978.
