## Supplementary material for "Body-plan reorganization in a sponge correlates with microbiome change": Vargasetal_Supplementary_Table_06

**Supplementary Table 6**. Top-50 Biological Process GO terms enriched among the set of transcripts differentially expressed in shaded *vs*. control *Lendenfeldia chondrodes* explants. Terms in boldface are related to developmental/morphogenetic processes or processes involving cell movement.

| GO term ID | Term | Annotated | Significant | Expected | classic |
|---|---|---|---|---|---|
| **GO:0031344** | **regulation of cell projection organization** | 69 | 33 | 19.15 | 0.00024 |
| **GO:0030595** | **leukocyte chemotaxis** | 15 | 11 | 4.16 | 0.00030 |
| GO:0050731 | positive regulation of peptidyl-tyrosine... | 13 | 10 | 3.61 | 0.00031 |
| **GO:0050920** | **regulation of chemotaxis** | 31 | 18 | 8.6 | 0.00034 |
| **GO:0006898** | **receptor-mediated endocytosis** | 60 | 29 | 16.65 | 0.00045 |
| **GO:0050900** | **leukocyte migration** | 39 | 21 | 10.82 | 0.00046 |
| **GO:0050767** | **regulation of neurogenesis** | 77 | 35 | 21.37 | 0.00052 |
| **GO:0009719** | **response to endogenous stimulus** | 130 | 53 | 36.08 | 0.00062 |
| GO:0051259 | protein oligomerization | 61 | 29 | 16.93 | 0.00064 |
| **GO:0021772** | **olfactory bulb development** | 16 | 11 | 4.44 | 0.00072 |
| **GO:0021988** | **olfactory lobe development** | 16 | 11 | 4.44 | 0.00072 |
| GO:0050730 | regulation of peptidyl-tyrosine phosphor... | 14 | 10 | 3.89 | 0.00081 |
| GO:0006887 | exocytosis | 51 | 25 | 14.15 | 0.00087 |
| **GO:0002688** | **regulation of leukocyte chemotaxis** | 10 | 8 | 2.78 | 0.00087 |
| GO:0043552 | positive regulation of phosphatidylinosi... | 10 | 8 | 2.78 | 0.00087 |
| GO:0090218 | positive regulation of lipid kinase acti... | 10 | 8 | 2.78 | 0.00087 |
| GO:1903727 | positive regulation of phospholipid meta... | 10 | 8 | 2.78 | 0.00087 |
| GO:0051289 | protein homotetramerization | 12 | 9 | 3.33 | 0.00088 |
| **GO:0030111** | **regulation of Wnt signaling pathway** | 38 | 20 | 10.55 | 0.00092 |
| **GO:0051240** | **positive regulation of multicellular organismal process** | 111 | 46 | 30.8 | 0.00094 |
| **GO:0006935** | **chemotaxis** | 100 | 42 | 27.75 | 0.00114 |
| **GO:0045664** | **regulation of neuron differentiation** | 66 | 30 | 18.32 | 0.00132 |
| **GO:0010975** | **regulation of neuron projection development** | 55 | 26 | 15.26 | 0.00136 |
| GO:1901652 | response to peptide | 34 | 18 | 9.44 | 0.00152 |

| | | | | | |
|---|---|---|---|---|---|
| GO:1901653 | cellular response to peptide | 29 | 16 | 8.05 | 0.00157 |
| **GO:0060284** | **regulation of cell development** | 87 | 37 | 24.14 | 0.00171 |
| **GO:0035567** | **non-canonical Wnt signaling pathway** | 22 | 13 | 6.11 | 0.00190 |
| **GO:0031346** | **positive regulation of cell projection organization** | 32 | 17 | 8.88 | 0.00196 |
| **GO:0060326** | **cell chemotaxis** | 27 | 15 | 7.49 | 0.00201 |
| **GO:0032755** | **positive regulation of interleukin-6 production** | 13 | 9 | 3.61 | 0.00215 |
| GO:0060401 | cytosolic calcium ion transport | 13 | 9 | 3.61 | 0.00215 |
| GO:0060402 | calcium ion transport into cytosol | 13 | 9 | 3.61 | 0.00215 |
| GO:1901700 | response to oxygen-containing compound | 130 | 51 | 36.08 | 0.00219 |
| **GO:0051960** | **regulation of nervous system development** | 94 | 39 | 26.09 | 0.00225 |
| **GO:0042330** | **taxis** | 103 | 42 | 28.58 | 0.00229 |
| **GO:0002685** | **regulation of leukocyte migration** | 11 | 8 | 3.05 | 0.00242 |
| **GO:0007405** | **neuroblast proliferation** | 11 | 8 | 3.05 | 0.00242 |
| **GO:0021889** | **olfactory bulb interneuron differentiation** | 11 | 8 | 3.05 | 0.00242 |
| **GO:0021891** | **olfactory bulb interneuron development** | 11 | 8 | 3.05 | 0.00242 |
| **GO:0032677** | **regulation of interleukin-8 production** | 11 | 8 | 3.05 | 0.00242 |
| **GO:0032757** | **positive regulation of interleukin-8 production** | 11 | 8 | 3.05 | 0.00242 |
| GO:0043550 | regulation of lipid kinase activity | 11 | 8 | 3.05 | 0.00242 |
| GO:0043551 | regulation of phosphatidylinositol 3-kin... | 11 | 8 | 3.05 | 0.00242 |
| GO:1903725 | regulation of phospholipid metabolic pro... | 11 | 8 | 3.05 | 0.00242 |
